# Nanoscale subcellular architecture revealed by multicolor 3D salvaged fluorescence imaging

**DOI:** 10.1101/613174

**Authors:** Yongdeng Zhang, Lena K. Schroeder, Mark D. Lessard, Phylicia Kidd, Jeeyun Chung, Yuanbin Song, Lorena Benedetti, Yiming Li, Jonas Ries, Pietro De Camilli, James E. Rothman, David Baddeley, Joerg Bewersdorf

## Abstract

Combining the molecular specificity of fluorescent probes with three-dimensional (3D) imaging at nanoscale resolution is critical for investigating the spatial organization and interactions of cellular organelles and protein complexes. We present a super-resolution light microscope that enables simultaneous multicolor imaging of whole mammalian cells at ~20 nm 3D resolution. We show its power for cell biology research with fluorescence images that resolved the highly convoluted Golgi apparatus and the close contacts between the endoplasmic reticulum and the plasma membrane, structures that have traditionally been the imaging realm of electron microscopy.

**One Sentence Summary:** Complex cellular structures previously only resolved by electron microscopy can now be imaged in multiple colors by 4Pi-SMS.

## Main Text

While ‘form follows function’ is a well-established principle in architecture, applying this concept to understand basic operating principles of a cell has been hampered by the lack of tools to resolve subcellular architecture. Revealing the intricate inner workings of cells requires visualizing the interactions between proteins and organelles with molecular specificity at nanoscale resolution in three dimensions (3D). The diffraction-limited resolution of conventional light microscopy (about 250 nm) stands in stark contrast to the structural dimensions of many organelles and complexes, such as the thickness of Golgi cisternae (about 50 nm each) (*1*) and the diameter of tubules of the endoplasmic reticulum (ER) (about 90 nm) (*2*). Electron microscopy (EM), while providing sufficient resolution with ease, lacks the capabilities of fluorescence light microscopy which offers excellent molecular specificity. Recently developed fluorescence super-resolution techniques have overcome the diffraction barrier and achieved impressive resolutions (*3, 4*). The ultimate goal, however - simultaneously resolving multiple targets of interest, for example the spatial relationship between two proteins of interest in the context of a related organelle in 3D - is still very challenging and has constrained the impact of super-resolution microscopy in cell biology.

To address this challenge, we set out to develop a super-resolution instrument which can obtain high-quality images in three color channels, i.e. 10-20 nm 3D resolution, high molecular detection efficiency and negligible channel shift and cross-talk. Two previous inventions in the super-resolution field form the foundation of our development: (i) interferometric detection of fluorescence from individual emitters by two opposing objectives in a ‘4Pi’ geometry with single-molecule switching (4Pi-SMS) has demonstrated nearly isotropic 3D resolution of ~10-20 nm (*5–8*). This imaging modality has shown excellent localization precision and densities of localized emitters in a single color channel, but has not been able to demonstrate similar image quality in additional color channels (*7–9*). (ii) Ratiometric color assignment can determine molecular identities based on the spectral information extracted from spectrally similar, simultaneously imaged fluorescent emitters (*10–14*). This approach allows for the use of dyes with the best optical switching properties which all emit in the far-red wavelength range (*15*) and avoids chromatic aberration problems and sequential imaging of dyes which is prone to drift and bleaching. Ratiometric color assignment has struggled so far, however, with obtaining high assignment efficiency without rejecting or falsely assigning large fractions of molecules, and doing so without substantially compromising localization precision. Combining interferometric 4Pi-SMS imaging with a refined ratiometric detection scheme which takes advantage of ‘salvaged fluorescence’ (**SF**), we show in this work high-localization density imaging at 10-20 nm 3D resolution in three color channels of whole mammalian cells.

Ratiometric single-molecule imaging assigns molecular identity by comparing the single-molecule emitter signal levels detected in two or more spectral windows (*10–14*). If emission spectra are known and signal to noise ratio is sufficiently high, two spectral windows are sufficient to distinguish more than two, in theory an arbitrarily large number of, different fluorescent probes (*10*). The classical implementation of ratiometric single-molecule imaging inserts a dichroic beamsplitter into the fluorescence beam path to create these two spectral detection windows. We realized that the main dichroic beamsplitter used in most fluorescent microscopes to separate the illumination from the fluorescence light already represents two spectral windows: the main transmitted, longer-wavelength component (conventional fluorescence) and a small but non-negligible reflected fraction (**Fig. 1A** and **fig. S1**). Salvaging this reflected fluorescence (salvaged fluorescence) provides previously lost spectral information which can be used to assign the molecular identity of an emitter. This approach takes advantage of the fact that spectral assignment and spatial localization precision utilize the fluorescent signal very differently. The former takes advantage of differences between probe spectra, which, given the steep rising edge of the emission spectra, are detected very clearly in the reflected spectral window. This suggests, that the salvaged fluorescence window can be quite narrow. The latter depends on the total photon number which, with a narrow salvaged fluorescence window, is mostly collected in the conventional fluorescence channel. Molecules can then be localized based on this channel alone avoiding the need for accurate registration and chromatic corrections necessary in classical ratiometric imaging which combines the signal of both channels. Simulations showed that with a transition edge between windows in the 660-670 nm range, dyes excitable at 642 nm and well suitable for SMS (*13*) can be separated very well (cross-talk fractions 1% to 2%, rejection fractions <1% to 10%) with only minor compromises (~1 nm) in localization precision (**fig. S2**).

**Fig. 1.**
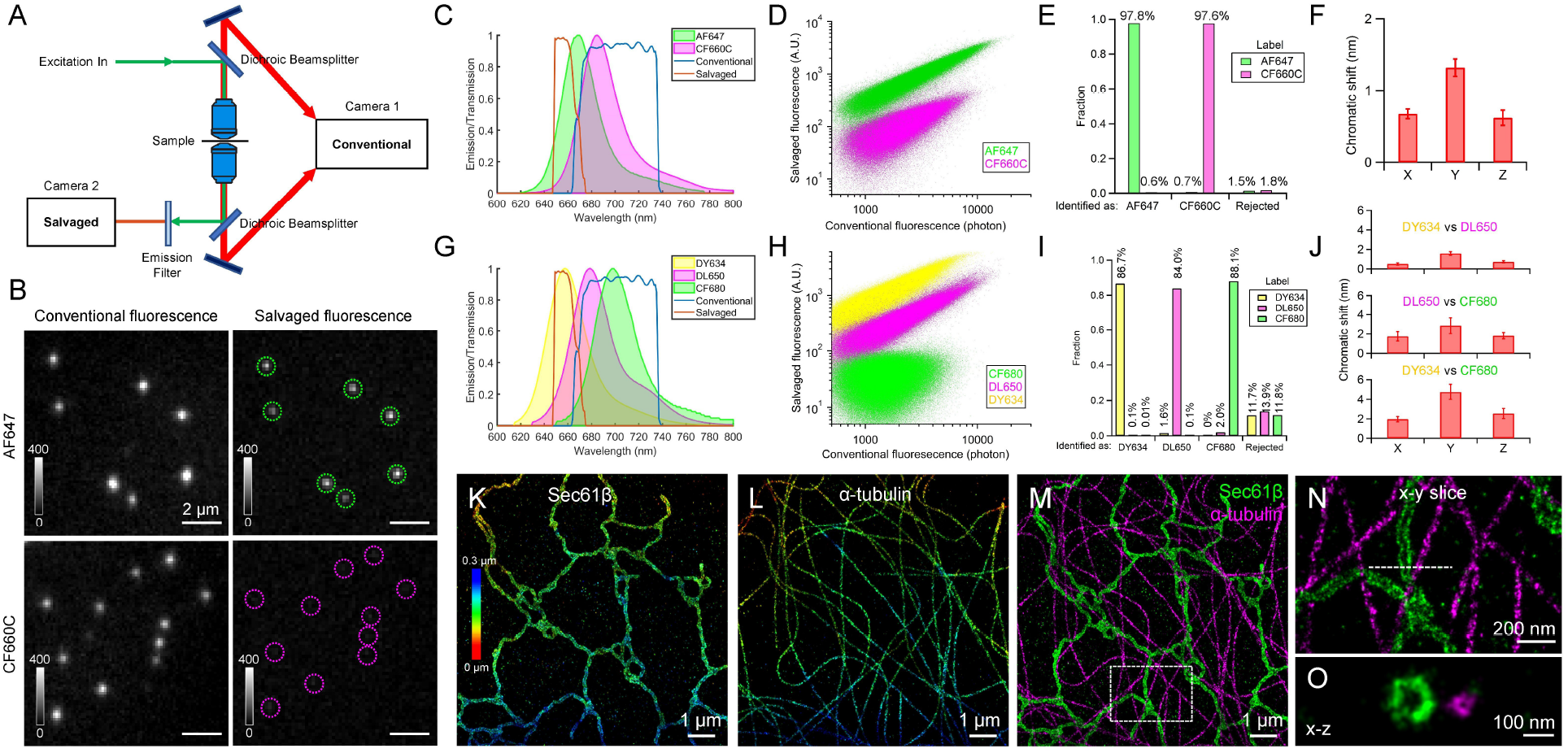
Characterization of multicolor 4Pi-SMS imaging using salvaged fluorescence. (**A**) Schematic of the multicolor 4Pi-SMS microscope. Camera 1 captures conventional fluorescence. Camera 2 captures salvaged fluorescence. (**B**) Conventional and salvaged fluorescence images of single molecules of AF647 and CF660C (Movie S1). Dashed circles indicate positions of single molecules observed in the conventional fluorescence channel. (**C**) Emission spectra of AF647 and CF660C and transmission profiles for conventional and salvaged fluorescence. (**D**) Scatter plot of salvaged fluorescence versus conventional fluorescence intensities of localized dye molecules in two single-color microtubule samples (shown in fig. S4) on a logarithmic scale. (**E**) Cross-talk and rejected fraction for the two dyes. (**F**) Chromatic shift between AF647 and CF660C in each dimension, determined from 23 images (each 20 x 20 μm). (**G**) Emission spectra of DY634, DL650 and CF680 and the transmission profiles for conventional and salvaged fluorescence. (**H**) Scatter plot of salvaged fluorescence versus conventional fluorescence intensities of localized dye molecules in three single-color microtubule samples (shown in fig. S5) on a logarithmic scale. (**I**) Cross-talk and rejected fraction for the three dyes. (**J**) Chromatic shift between each dye pair, determined from 13 images (each 20 x 20 μm). (**K** to **M**) Two-color images of microtubules and ER membrane in a COS-7 cell. (**N**) 50-nm thick x-y slice of the boxed region in (M). (**O**) 20-nm thick x-z cross-section along the dashed line in (N). Data are presented as mean ± SEM in (**F** and **J**).

We implemented the SF approach in a 4Pi-SMS instrument (**Fig. 1A** and **fig. S1**) and tested it with five far-red dyes (Alexa Fluor 647 (AF647), CF660C, Dyomics 634 (DY634), Dylight 650 (DL650), CF680) for two- and three-color imaging (**Fig. 1** and **fig. S3**). For two-color imaging with AF647 and CF660C (Fig. 1, B to F; **Movie S1**), we achieved a localization precision (~7 nm in xy, ~5 nm in z) (**fig. S4, A** to **F**) that is comparable to the previously reported one-color 4Pi-SMS imaging (*8*). Consistent with these high localization precision values, the microscope can resolve the hollow center of labeled ER tubules and microtubules (Fig. 1, K to O). The distinct ratios of salvaged to conventional fluorescence between the two dyes yielded a cross-talk of < 1% (Fig. 1, D and E; **fig. S4, G** and **H**). The chromatic shifts between the two channels were determined to be less than 1.5 nm in all directions without applying any chromatic corrections (**Fig. 1F**; **fig. S4, I** to **K**).

Using DY634, DL650 and CF680 for three-color imaging (Fig. 1, G to J), we resolved the tubular structure of immunolabeled microtubules in all three color channels (**fig. S5, A** to **I**), and achieved ≤ 2% cross-talk between each dye pair (**Fig. 1I**; **fig. S5, J** and **K**). The chromatic shifts between the three dyes were determined to be mostly in the 1-3 nm range with the biggest value being < 5 nm for the dyes with the farthest shifted spectra (**Fig. 1J**; **fig. S5, L** and **M**).

We further tested our SF approach by imaging spatially-close cellular structures with known geometries (**Fig. 2**). The luminal and membrane markers of ER were resolved with no apparent chromatic aberrations despite the small diameter (~90 nm) (Fig. 2, A to E; **fig. S6, A** to **C**; **Movie S2**). The high 3D resolution revealed the subtle differences between two immunolabeling approaches: the apparent ER tubule diameter is about 20 nm smaller when labeled with a nanobody compared to conventional primary and secondary antibodies (**fig. S6, D** to **I**), which is consistent with their expected sizes (*16*). Mitochondrial nucleoid DNA was positioned within and clearly isolated from the boundaries of the outer mitochondrial membrane at various depths (Fig. 2, F to I; **fig. S7, A** to **C**; **Movie S3**). In contrast, two outer mitochondrial membrane proteins were well colocalized throughout the volume of the sample (**fig. S7, D** to **K**), confirming that our method results in precise alignment of structures without applying any chromatic corrections, even in thicker volumes.

**Fig. 2.**
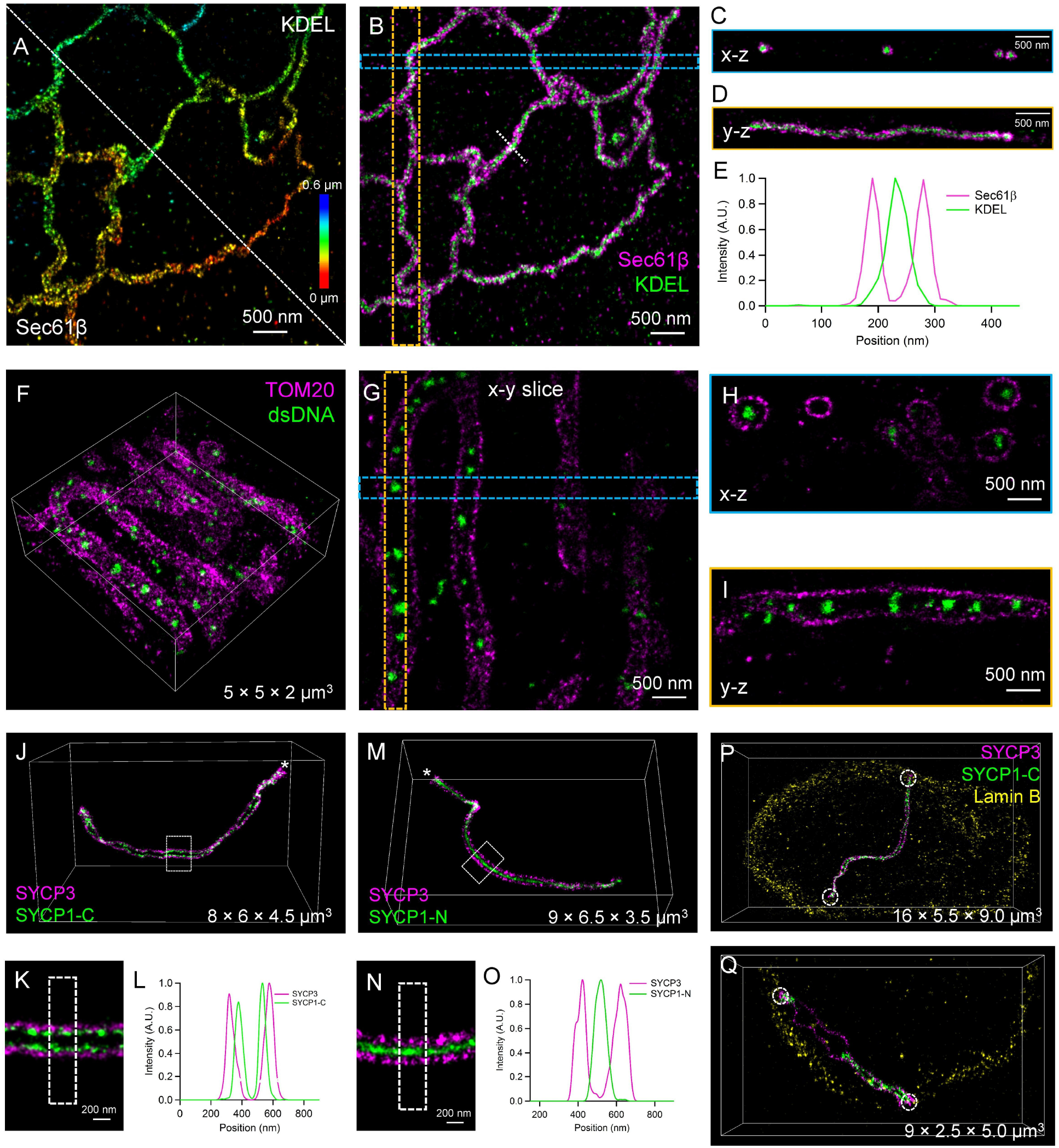
Multicolor 4Pi-SMS images of ER, mitochondria, and synaptonemal complexes. (**A** and **B**) Two-color image of ER membrane and lumen in a COS-7 cell (Movie S2). Rainbow color denotes depth in (A). Overlay image in (B). (**C**) x-z view of the blue dashed box (200 nm wide) in (B). (**D**) y-z view of the orange dashed box (400 nm wide) in (B). (**E**) Intensity profile along the white dashed line in (B). (**F**) Two-color image of outer mitochondrial membrane and mitochondrial DNA in a HeLa cell (Movie S3). (**G**) A 200-nm thick x-y slice of the image in (F). (**H**) x-z view of the blue dashed box (300 nm wide) in (G). (**I**) y-z view of the orange dashed box (300 nm wide) in (G). (**J**) Computationally isolated two-color image of SYCP3 and SYCP1-C in a synaptonemal complex (fig. S8, A and B; Movie S4, part I). (**K**) Magnified image of the boxed region in (J). (**L**) Intensity profile along the dashed boxed region in (K). (**M**) Computationally isolated two-color image of SYCP3 and SYCP1-N in a synaptonemal complex (fig. S8, C and D; Movie S4, part II). (N) Magnified image of the boxed region in (M). (**O**) Intensity profile along the dashed boxed region in (N). (**P** and **Q**) Three-color images of SYCP3 and SYCP1-C on two computationally isolated synaptonemal complexes next to Lamin B (fig. S8E; Movie S4, part III). White dashed circles indicate where synaptonemal complexes contact the nuclear lamina.

To demonstrate the power of the SF approach in thicker cells, we imaged the synaptonemal complex (SC) in intact mouse spermatocytes (Fig. 2, J to Q; **fig. S8**; **Movie S4**). Our method revealed the twisted helical structure of two synaptonemal scaffold proteins, SYCP1 and SYCP3, throughout the 10-μm thick volume (Fig. 2, J to O; **fig. S8, A** to **D**). Furthermore, imagining N- and C-terminally labeled SYCP1 confirmed that the C-terminus is oriented towards the SYCP3 tracks while the N-terminus extends into the central region (Fig. 2, J and M) (*17*). Imaging these two proteins alongside Lamin B showed the ends of the SCs connecting to the nuclear lamina (**fig. S8, E** and **F**), confirming previous results (*18*). The ends of fully assembled (**Fig. 2P**) as well as partially assembled SCs (**Fig. 2Q**) are closely associated with the lamina.

The minimal cross-talk and negligible chromatic aberrations make the SF approach a powerful tool for cell biology. To test how well it can reveal the location of proteins in complex 3D morphologies that are otherwise only accessible through EM, we imaged two challenging structures which are of central importance to cell biology: the Golgi apparatus and contact sites between the ER and the plasma membrane (PM).

We imaged the immunolabeled Golgi apparatus in HeLa cells in three colors (Fig. 3, A to E; **fig. S9**; **Movie S5**). The *cis, medial*, and *trans* regions appeared as distinct structures stacked parallel to each other in cross-sections through the imaged Golgi apparatus (Fig. 3, F and G). While the *cis-medial-trans* stacking was maintained throughout the Golgi, some stacks flipped their orientation within a few hundred nanometers (**Fig. 3F**, compare dashed line and cyan arrow). The high 3D resolution enabled us to characterize the distance between *cis*, *medial* and *trans* cisternae despite the convoluted morphology of the Golgi apparatus (Fig. 3, H and I). The *trans*-localized p230 and *cis*-localized GRASP65 labels showed a peak-to-peak distance of on average 187 ± 6 nm (mean ± SEM). The broad distribution of the peak-to-peak distances appeared to consist of two populations which suggests that we observed Golgi stacks with different numbers of cisternae. The *medial*-localized ManII was on average 85 ± 3 nm (mean ± SEM) apart from GRASP65. In contrast, GRASP65 and the also *cis*-localized GM130 showed an average separation not significantly different from zero (**fig. S10**), confirming previous biochemical data (*19*). It has been previously reported that each Golgi cisterna is about 50 nm thick (*1*). Side profile measurements of our data showed that both p230 and GM130 stainings had a thickness of about 70-80 nm (Fig. 3, H and J). This is an overestimate as we averaged across 1 μm subregions and did not account for the fact that the cisternae are not perfectly flat over these regions. Considering the size of primary and secondary antibodies (> 20 nm combined; **fig. S6I**), this data suggests, that p230 and GM130 are enriched in one cisterna only, as expected. Our GRASP65 staining showed a similar thickness as p230 and GM130 (**Fig. 3J**), indicating that our GRASP65 labeling is concentrated in one cisterna. In contrast, ManII-GFP appeared at a thickness of about 120-130 nm implying that it is distributed over multiple cisternae (**Fig. 3J**).

**Fig. 3.**
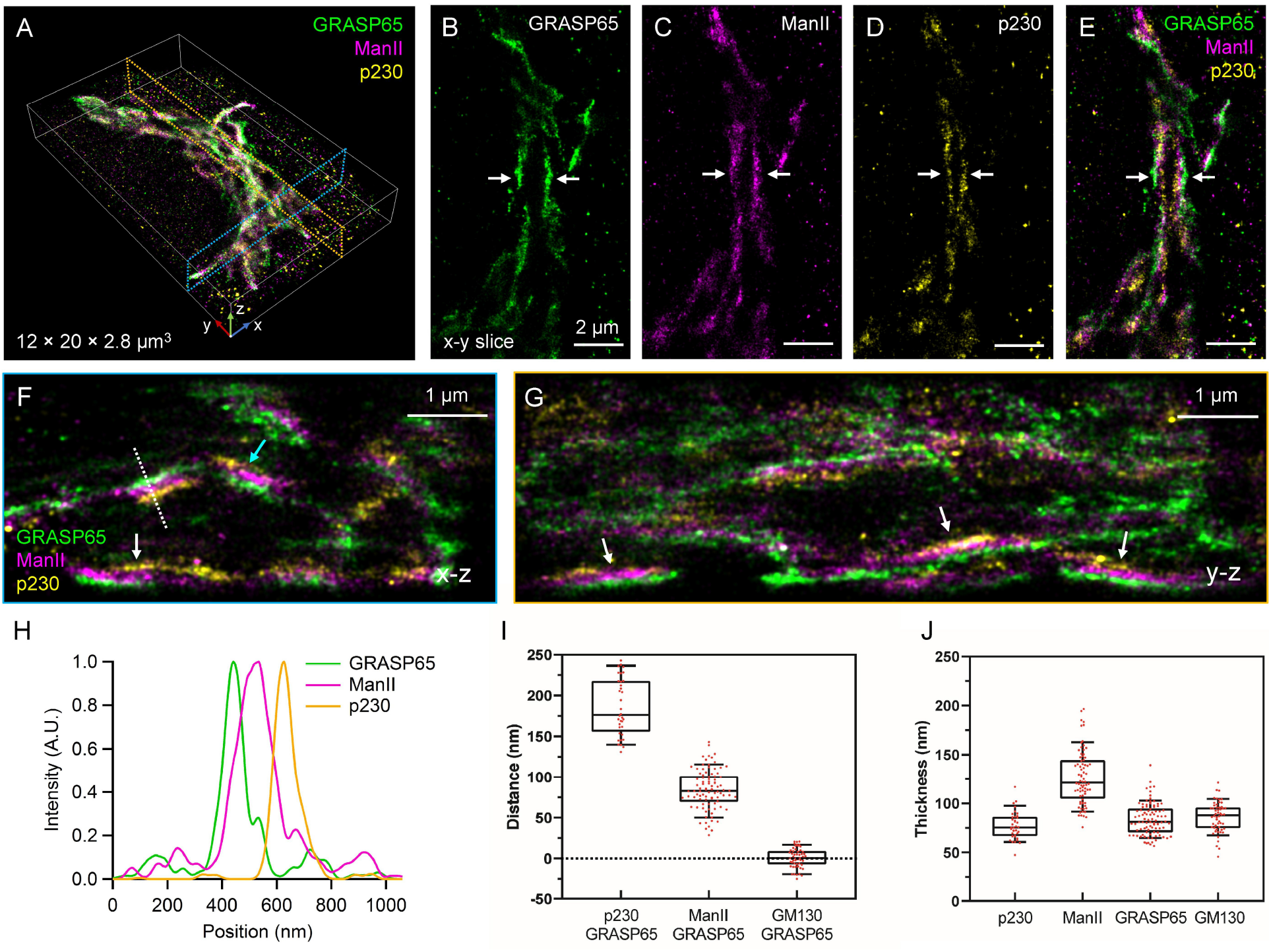
Complex stacked architecture of the Golgi apparatus. (**A**) 3D overlay image of *cis* (GRASP65), *medial* (ManII), and *trans* (p230) Golgi proteins in a HeLa cell (fig. S9; Movie S5, part I). (**B** to **E**) 500-nm thick x-y slice of data shown in (A). (**F**) 1-μm thick x-z cross-section centered at the blue dashed region in (A). (**G**) 1-μm thick y-z cross-section centered at the orange dashed region in (A). (**H**) Intensity profile along the dashed line in (F). (**I**) Distance between the peak intensities of each pair of Golgi proteins (p230-GRASP65, n= 36 from 9 cells; ManII-GRASP65, n = 90 from 18 cells; GM130-GRASP65, n = 54 from 9 cells). (**J**) Thickness of Golgi regions, measured as the full-width at half-maximum of the intensity profile (p230, n = 36 from 9 cells; ManII, n = 90 from 18 cells; GRASP65, n = 90 from 18 cells; GM130, n = 54 from 9 cells). Arrows denote where the *cis-medial-trans* stack is visible (B to E, F and G). Median and interquartile range are shown with whiskers drawn down to the 10th percentile and up to the 90th percentile (I and J).

Unlike the perinuclear Golgi, the ER spreads and branches throughout the volume of mammalian cells making contacts with most organelles, including the PM. To gain insight into fine details of ER-PM contacts, where the intermembrane distance is typically within the 15-25 nm range (*20*), we imaged the PM (labeled with WGA) and ER membranes in COS-7 cells (**Fig. 4**; **Movie S6**). At the periphery of the cells, ER tubules are generally clearly separated by tens of nanometers from the top and bottom PM (**Fig. 4A**). Upon overexpression of an ER protein, ORP5, which functions as a tether at ER-PM contact sites, however, a large fraction of the peripheral ER became closely apposed to the PM to form large, patch-like contacts (**Fig. 4B**, cyan arrow) (*21*). Likewise, overexpression of another ER-PM contact site protein, ESYT2 (*22*), also expanded appositions of ER with the PM, although in this case, the ER retained a tubular shape at such appositions (**fig. S11A**). This was consistent with ESYT2 being anchored to the ER membrane by an N-terminal hairpin domain that may sense/induce high-curvature membranes (*22*), while ORP5 is anchored to the ER by a single C-terminal transmembrane region (*21*). To visualize the contact site proteins directly, we imaged ORP5 or ESYT2 together with the PM marker (Fig. 4, C to F; **fig. S11, B** and **C**). In agreement with the results shown above (*21*), ORP5 appeared as patches, while ESYT2 appeared as tubule-like structures (**fig. S11, D** to **F**). Two-color imaging further confirmed that the observed ORP5 and ESYT2 structures corresponded to ER elements (**fig. S12, A** to **F**). The intensity profile of each cross-section showed a small separation between ORP5 or ESYT2 and the PM (Fig. 4, D and F; **fig. S11F**, blue arrow). Quantification showed a distance of the PM signal peak to the ORP5 signal peak of 17 ± 9 nm (mean ± SD), while the distance from the ESYT2 signal peak was slightly greater, 20 ± 11 nm (mean ± SD) (**Fig. 4G**), consistent with values derived from EM images (*20*) and demonstrating the power of SF in resolving ultrastructural details. While it is difficult to visualize specific contact site proteins in EM, our approach allows imaging the ER membrane, contact site proteins, and the PM at the same time. Three-color imaging of ER-PM contact sites showed both contact site proteins as expected at the interface between ER membranes and the PM: ORP5 at patch-like contacts (Fig. 4, H and I; **fig. S12G**) and ESYT2 at tubular contacts (Fig. 4, J and K; **fig. S12H**). In addition, both ORP5 and ESYT2 localized to the ER membrane facing the PM but not the other side (Fig. 4, H to K; **Movie S7**).

**Fig. 4.**
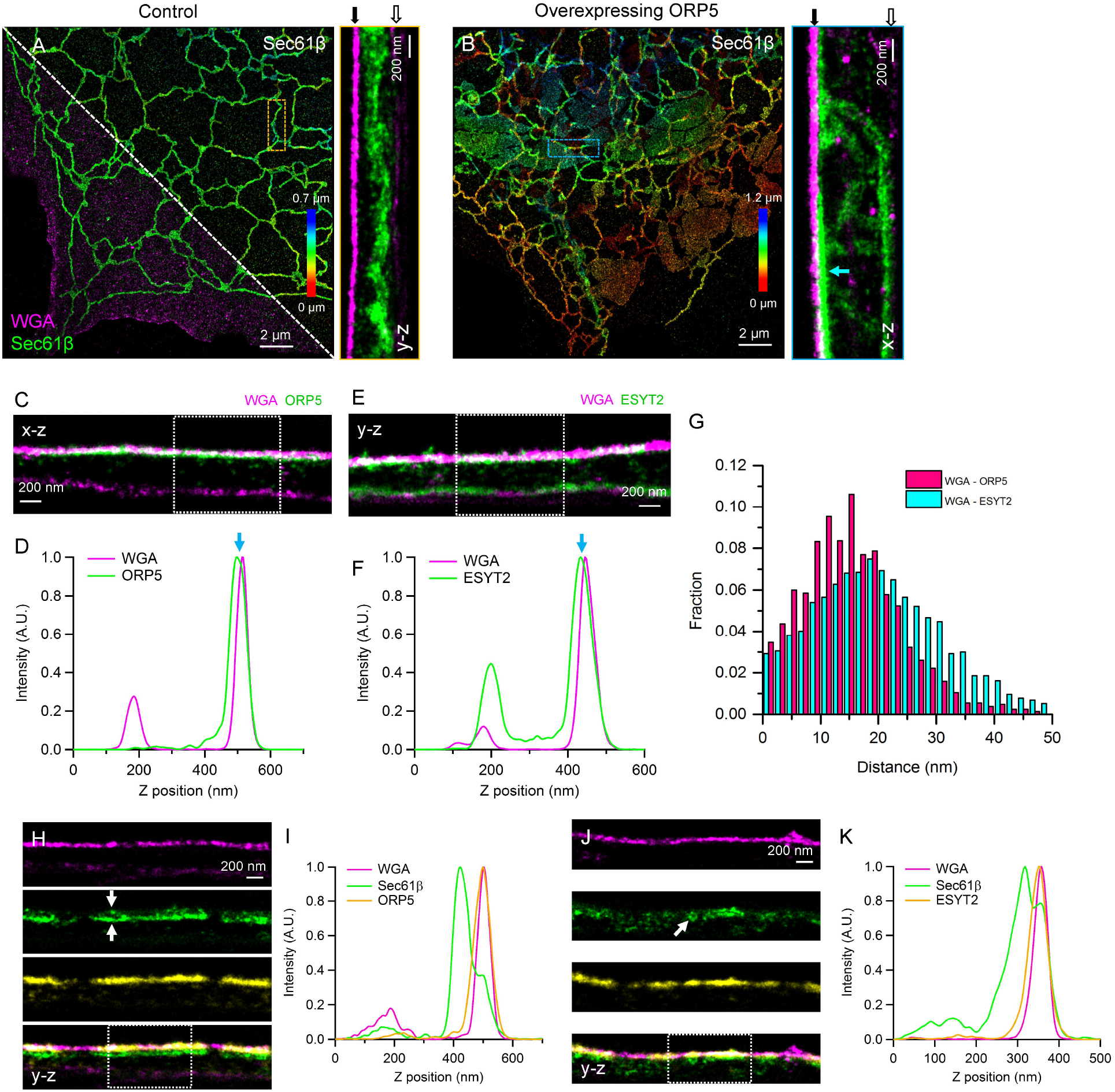
The 3D architecture of ER-PM contact sites. (**A**) Two-color image of ER (Sec61β) and PM (WGA) in a COS-7 cell (Movie S6, part I). Top right half shows the ER where rainbow color denotes depth. Bottom left half overlays ER and PM. Right panel shows magnified y-z view of boxed region. (**B**) Two-color image of ER and PM in a cell overexpressing mCherry-ORP5 (Movie S6, part II). Right panel shows magnified x-z view of boxed region. Solid and outlined arrows in (A and B) point to the top and bottom PM, respectively. (**C**) Two-color ORP5 and PM x-z view, overview shown in (fig. S11B). (**D**) Axial intensity profile across the dashed box in (C). (**E**) Two-color ESYT2 and PM y-z view, overview shown in (fig. S11C). (**F**) Axial intensity profile across the dashed box in (E). Blue arrows in (D and F) indicate the distance between PM and contact site proteins. (**G**) Histograms of the distance between PM and contact site proteins (from > 2800 subregions of 100 x 100 nm size, n = 4 cells per condition) (See Material and Methods). (**H**) Three-color ER, ORP5 and PM y-z view, overview shown in (fig. S12G) (Movie S7, part I). White arrows indicate the top and bottom membranes of the ER. (I) Axial intensity profile across the dashed box in (H). (**J**) Three-color ER, ESYT2, and PM y-z view, overview shown in (fig. S12H) (Movie S7, part II). White arrow points to an ER tubule. (**K**) Axial intensity profile across the dashed box in (J).

It is important to note that our SF approach can also be implemented in single-objective systems with minor optical modifications (**fig. S13, A** and **B**). To show the feasibility, we blocked the top emission beam path in our 4Pi-SMS microscope to mimic the detection of a single-objective system (**fig. S13C**). We obtained two-color images with excellent quality in both 2D and 3D with astigmatism (**fig. S13, D** to **J**). Additionally, the concept can be adopted in many multicolor imaging scenarios, including other single-molecule imaging techniques and single-particle tracking.

The development of high-quality multicolor imaging at tens of nanometers resolution in 3D provides a tool to the cell biologist that combines the strength of specific labeling of fluorescence microscopy in the context of interaction partners and cellular landmarks with a level of detail that traditionally had been the realm of EM.

## ACKNOWLEDGMENTS

We would like to thank Ricardo Benavente (University of Wuerzburg) for sharing the anti-SYCP1-N antibody, Andrew Barentine and Andreas Ernst for comments on the manuscript, and Carl Ebeling (Bruker Corp.) for help with the Vutara SRX software.

## Funding

This work was primarily supported by grants from the Wellcome Trust (203285/B/16/Z), the G. Harold and Leila Y. Mathers Foundation, and NIH (R01 GM118486, P30 DK045735 and NS36251). Y.L. acknowledges support from the EMBL Interdisciplinary Postdoc Programme (EIPOD) under Marie Curie Actions COFUND.

## Author contributions

Y.Z. and J.B. conceived the idea. Y.Z. implemented the hardware. Y.Z., Y.L., J.R., and D.B. wrote the software. L.K.S., J.C., P.D.C., J.E.R., and J.B. designed biological experiments. L.K.S. developed sample preparation protocols. L.K.S. prepared and optimized microtubule, ER, mitochondria, Golgi, and ER-PM contact sites samples. Y.S. and L.B. provided the synaptonemal complex samples. M.L. prepared and optimized the synaptonemal complex samples. P.K. prepared and optimized microtubule samples. Y.Z. imaged the samples and analyzed the images. Y.Z., L.K.S., and J.B. wrote the manuscript with input from all authors. All authors discussed the results and commented on the manuscript.

## Competing interests

J.B. has financial interests in Bruker Corp. and Hamamatsu Photonics. J.B. is co-inventor of a U.S. patent application (US20170251191A1) related to the 4Pi-SMS system and image analysis used in this work. Y.Z. and J.B. have filed a U.S. provisional patent application (No. 62/723,291) about the salvaged fluorescence multicolor imaging method described in this work.

## SUPPLEMENTAL MATERIALS

Materials and Methods

Figs. S1 to S13

References (*23-28*)

Movies S1 to S7

