## Supplementary Material for "Nanoscale subcellular architecture revealed by multicolor 3D salvaged fluorescence imaging"

**This PDF file includes:**

Materials and Methods
Figs. S1 to S13
References (1-28)

**Other Supplementary Materials for this manuscript include the following:**

Movies S1 to S7

### Materials and Methods

#### Synaptonemal complex samples

All experimental procedures involving the use of mice were performed in agreement with the Yale University Institutional Animal Care and Use Committee (IACUC). BALB/cJ mice (The Jackson Laboratory, Stock No: 000651) were purchased from The Jackson Laboratory. Testes (tunica removed) from 18-day old mice were disrupted using forceps and a razor blade in 1 mL of PBS (1× PBS; Gibco, Cat# 10010023) with protease inhibitors (Roche, Complete Ultra, Cat# 05896988001). The cell suspension was then gently added to a 15-mL conical tube with 5 mL of 1× PBS with protease inhibitors and allowed to settle. After approximately 3 minutes, 5 1-mL aliquots of the cell suspension were placed in 1.5-mL microcentrifuge tubes and centrifuged at 9k RPM for 10 minutes. The supernatant was then aspirated, and the pellets were combined in 0.5 mL of 1× DPBS (made from 10x stock; Gibco, Cat#14080-055) per testes. 50-100 µL of the cell suspension was added to #1.5, 25-mm diameter round precision coverglass, which had been cleaned and Poly-L-Lysine coated (Sigma-Aldrich, Cat# P4707), and allowed to sit for 30 minutes. The cells on coverglass were then fixed in 4% paraformaldehyde (PFA; Electron Microscopy Sciences, Cat# 15710) for 15 minutes at room temperature. Standard immunolabeling was performed using the antibodies listed in the Key Resource Table (See synaptonemal complex labeling section).

#### Cell culture

COS-7 and HeLa cells were grown in DMEM or DMEM/F12 (Gibco, Cat# 21063029 and 21041025) supplemented with 10% fetal bovine serum (Gibco, Cat# 10438026) at 37 °C with 5% $CO_2$. Some cultures were grown with media supplemented with sodium pyruvate (Gibco, Cat# 11360070).

#### Plasmids

For labeling the ER membrane, we expressed mEmerald-Sec61-C-18, a gift from Michael Davidson (deceased, formerly Florida State University, Tallahassee, FL; Addgene plasmid # 54249; referred to as GFP-Sec61β). For labeling mitochondria, we expressed GFP-OMP25 which was made by modifying pEGFP-C1 (Takara Bio Inc.) to include eGFP fused to the C-terminus of human OMP25/SYNJ2BP which contains the amino acids "QVQNGPIGHRGEGDPSGIPIFMVLVPVFALTMVAAWAFMRYRQQL" and localizes to mitochondria, as shown previously (*23*). For labeling the *medial* Golgi, we expressed GFP-ManII which was made from pEGFP-N1 (Takara Bio Inc.) to include amino acids 1-137 of mouse MAN2A1 fused to GFP, such that GFP is located in the Golgi lumen. For labeling the ER lumen, we expressed mCherry-KDEL. This was made by modifying pDsRed2-ER (Takara Bio Inc.), which encodes a signal peptide fused to the DsRed2 fluorescent protein followed by the tetrapeptide ER retention signal KDEL. The mCherry gene was amplified using the following primers 5'-ATACCGGTCGATGGTGAGCAAGGGCGAG-3' and 5'-CTGAAGCTTTTACAGCTCGTCCTTCTTGTACAGCTCGTCCATGCC-3'. The pDsRed2-ER plasmid and the mCherry PCR product were both digested using AgeI and HindIII (New England Biolabs, Cat# R0552S and R3104S) and ligated together, replacing the DsRed gene with the mCherry gene and resulting in the mCherry-KDEL plasmid.

#### Coverglass cleaning

Precision thickness coverglass (Bioscience tools, Cat# CSHP-No1.5-25) was cleaned before cells were plated on them. Before plating COS-7 cells, glass was cleaned in a sonic bath (Bronson) immersed in 1M KOH for 15 minutes and then rinsed with MilliQ water three times. Glass was then sterilized with 100% Ethanol, incubated with Poly-L-Lysine for 10 minutes, then rinsed with sterile PBS before adding media and cells. Before plating HeLa cells, glass was instead cleaned with an ozone cleaner (Jelight, UVO Cleaner #342A) for 30 minutes. Media and HeLa cells were placed directly on ozone-cleaned glass. Before placing spermatocytes, glass was cleaned in a plasma oven for 5 minutes before being coated with Poly-L-Lysine.

#### Transfection

Samples including GFP-Sec61β, mCherry-KDEL and GFP-OMP25 expression in HeLa or COS-7 cells used DNA transfection by electroporation. DNA was introduced to HeLa or COS-7 cells using a NEPA GENE electroporation device. Approximately 1 million cells were rinsed in Opti-MEM (Gibco, 31985070) and then resuspend in Opti-MEM with 10 µg DNA in an electroporation cuvette with 2-mm gap (Bulldog Bio, Cat# 12358346). Cells were electroporated with a poring pulse of 125 V, 3-ms pulse length, 50-ms pulse interval, 2 pulses, with decay rate of 10% and + polarity; followed by a transfer pulse of 25 V, 50-ms pulse length, 50-ms pulse interval, 5 pulses, with a decay rate of 40% and ± polarity. After electroporation, cells were removed from the cuvette and grown in growth media on prepared coverglass. Samples were fixed 18-24 hours after electroporation (see coverglass cleaning section).

Samples including ER-PM contact site protein expression in COS-7 cells utilized Lipofectamine2000 (Invitrogen, Cat# 11668027) transfection. The day before transfection, cells were plated on prepared coverglass (see coverglass cleaning section). Transfection was performed as suggested by the manufacturer with a modification of 1 µL transfection reagent and 1 µg DNA per well in a 6-well plate. When two DNAs were transformed together, 0.5 µg of each DNA was used for a total of 1 µg DNA.

#### Fluorescent Dyes

Alexa Fluor 647 (AF647) and CF660C were used for all two-color imaging experiments except for the ER-PM imaging where CF680 was used instead of CF660C (see ER-PM contact sites labeling). Secondary antibodies labeled with AF647 (Invitrogen, Cat# A21236, A21237, or A21245; used at 1:1000 dilution for 1 hour at room temperature) and CF660C (Biotium, Cat# 20812-500µL or 20813-500µL; used at 1:500 or 1:1000 dilution for 1 hour at room temperature) were used to label primary antibodies. These CF660C-labeled antibodies were manufactured to have one dye per antibody. Nanobody GFP-binding protein (Chromotek, Cat# gt-250) and RFP-binding protein (Chromotek, Cat# rt-250) were conjugated to AF647 in lab (see nanobody and antibody conjugation section).

Dyomics634 (DY634), DyLight 650 (DL650), and CF680 were used for all three-color imaging experiments. Secondary antibodies labeled with DY634 (conjugated in lab, see nanobody and antibody conjugation section; used at 1:200 dilution for 1 hour at room temperature), DL650 (Invitrogen, Cat# SA5-10174 or SA5-10034; used at 1:1000 dilution for 1 hour at room temperature), and CF680 (Biotium, Cat# 20817-500µL or 20818-500µL; used at 1:500 or 1:1000 dilution for 1 hour at room temperature) were used to label primary antibodies. The CF680-labeled antibodies were manufactured to have one dye per antibody. Nanobody RFP-binding protein conjugated to DY634 and nanobody GFP-binding protein conjugated to DL650 were made in lab (see nanobody and antibody conjugation section). WGA-CF680 (Biotium, Cat# 29029-1) was used for plasma membrane labeling.

#### Nanobody and antibody conjugation

Nanobody RFP-binding protein or GFP-binding protein was conjugated to Alexa Fluor 647 NHS-ester (Life Technologies, Cat# A20006), DY634 NHS-ester (Dyomics, Cat# DY-634-NHS-ester portionized, 634-01A), or DL650 NHS-ester (Thermo Scientific, Cat# 62265). Approximately 100 µL conjugations were performed in 0.1 M Sodium Bicarbonate for 1 hour in the dark. Excess dye was removed from the conjugation reaction using Zeba Spin Desalting Columns with a 7K molecular weight cut off (Thermo Scientific, Cat# 89882). Nanobodies were used at 1:1000 dilution at room temperature or overnight at 4 °C.

Unlabeled goat anti-rabbit IgG, and goat anti-mouse IgG, and goat anti-human IgG (Jackson ImmunoResearch, Cat# 111-005-144, 115-005-146, and 109-005-088, respectively) were conjugated with DY634 NHS ester (Dyomics, DY-634-NHS-ester portionized, 634-01A). Approximately 100 µL reactions were performed in 0.1 M Sodium Bicarbonate for 1 hour in the dark. Excess dye was removed from antibody using Pro-Spin columns (Princeton Separations, Cat# CS800) per manufacturer's recommendations. Secondary antibodies conjugated in this work were used at 1:200 for 1 hour at room temperature.

#### Sample labeling

**Microtubule Samples.** To label microtubules, COS-7 cells were prepared as previously reported (*8*). After growing on Poly-L-Lysine coated coverglass for 24 hours, cells were rinsed with PBS, warmed to 37 °C , then incubated for 1 minute in 0.05% saponin diluted in cytoskeletal buffer (**CBS;** 10 mM MES pH 6.1, 138 mM NaCl, 3 mM MgCl2, 2 mM EGTA, 320 mM sucrose), warmed to 37 °C. Cells were subsequently fixed in 3% paraformaldehyde and 0.1% glutaraldehyde (GA; Electron Microscopy Sciences, Cat# 16019) diluted in CBS, warmed to 37 °C. Cells were permeabilized and blocked with 3% bovine serum albumin (BSA; Jackson ImmunoResearch, Cat# 001-000-162) and 0.2% Triton X-100 (TX-100; Sigma-Aldrich, Cat# T8787) in 1× PBS (diluted from 10× PBS; American Bio, Cat# AB11072-0100) for 30 minutes. The samples were incubated overnight at 4 °C with mouse anti-α-tubulin antibody (Sigma-Aldrich, Cat# T5168) diluted to 1:200 in antibody dilution buffer (1% BSA and 0.2% TX-100 in 1× PBS). Cells were washed in wash buffer (0.05% TX-100 in 1× PBS) three times for 5 minutes each. For single-color labeling, microtubule samples were incubated with each dye-conjugated secondary antibody (used at 1:1000) in antibody dilution buffer, for 1 hour at room temperature then washed three times for 5 minutes with wash buffer. For two- or three-color labeling, microtubule samples were incubated with the dye-conjugated secondary antibodies (used at 1:1000) together. Lastly, the samples were post-fixed in 3% PFA + 0.1% GA in CBS for 10 min, rinsed with PBS three times, and stored in PBS at 4 °C until imaged.

**ER and microtubule labeling.** COS-7 cells were electroporated with GFP-Sec61β plasmid and grown on Poly-L-Lysine coated glass for 24 hours before being fixed with 3% PFA + 0.1% GA in PBS for 15 minutes at room temperature. Samples were rinsed three times in 1× PBS before permeabilizing for 3 minutes using permeabilization buffer (**PB**; 0.3% CA-630 (Sigma-Aldrich, Cat# I8896), 0.05% TX-100, 0.1% BSA, and 1× PBS) at room temperature. Samples were then rinsed three more times with 1× PBS followed by 1 hour in block buffer (**BB**; 0.05% CA-630, 0.05% TX-100, 5% normal goat serum (Jackson ImmunoResearch, Cat# 005-000-121), and 1× PBS). Primary antibodies, rabbit anti-GFP (Invitrogen, Cat# A11122, used at 1:500) and mouse anti-tubulin (used at 1:1000), were diluted in BB and incubated with samples overnight at 4 °C. Samples were then washed in wash buffer (**WB**; 0.05% CA-630, 0.05% TX-100, 0.2% BSA, and 1× PBS) three times for 5 minutes each before secondary antibody labeling for 1 hour at room temperature diluted in BB. Samples were washed in WB three times for 5 minutes each. Post-fixation was performed using 3% PFA + 0.1% GA in 1× PBS for 10 minutes. Samples were rinsed three times in 1× PBS before being stored in 1× PBS at 4 °C.

**ER membrane and ER lumen labeling.** COS-7 cells were electroporated with GFP-Sec61β alone or GFP-Sec61β and mCherry-KDEL together. Cells were then transferred to cleaned and Poly-L-Lysine coated glass and grown for 24 hours before being fixed in 3% PFA + 0.1% GA in 1× PBS for 15 minutes.

All ER samples were permeabilized with **PB** (0.3% CA-630, 0.05% TX-100, 0.1% BSA, and 1× PBS) for 3 minutes at room temperature, rinse in 1× PBS three times, and then blocked in **BB** (0.05% CA-630, 0.05% TX-100, 5% normal goat serum, and 1× PBS) for an hour. Samples were washed in **WB** (0.05% CA-630, 0.05% TX-100, 0.2% BSA, and 1× PBS) three times for 5 minutes each after labeling with primary antibody, secondary antibody, or nanobody. The ER was labeled using different combinations of antibodies and/or nanobodies.

For two-color ER membrane labeling with antibodies, GFP-Sec61β was labeled with rabbit anti-GFP which was then labeled with two competing secondary antibodies. For two-color ER membrane labeling combining nanobody and antibody, GFP-binding protein nanobody, conjugated with AF647, first was used to label GFP-Sec61β at room temperature for 1 or 2 hours. Then rabbit anti-GFP (used at 1:500) was incubated with samples at 4 °C overnight, which was subsequently labeled with CF660C-ST. Two-color ER membrane and lumen labeling was performed on cells expressing GFP-Sec61β and mCherry-KDEL. Lumen-localized mCherry-KDEL was first labeled using RFP-binding protein nanobody conjugated to AF647 for 2 hours at room temperature. GFP-Sec61β was then labeled using rabbit anti-GFP (used at 1:500 at 4 °C overnight) which was then labeled with CF660C-ST. All samples were post-fixed using 3% PFA + 0.1% GA in 1× PBS for 10 minutes. Samples were rinsed three times in 1× PBS before being stored in 1× PBS at 4 °C.

**Mitochondria labeling.** HeLa cells were used for imaging mitochondria. For some samples, GFP-OMP25 plasmid was introduced to cells using electroporation and allowed to express for 18-24 hours. Mitochondria samples were fixed differently depending on the primary antibody being used, with a preference for 3% PFA + 0.1% GA in 1× PBS for 15 minutes at room temperature. Samples that included mouse anti-dsDNA antibody labeling, since this primary antibody does not label cells if GA is used during fixation, were fixed using 4% PFA in 1× PBS for 1 hour at room temperature. All cells were permeabilized with **PB** (0.3% CA-630, 0.05% Triton X-100, 0.1% BSA, and 1× PBS) for 3 minutes at room temperature, rinsed three times in PBS, and then blocked in **BB** (0.05% CA-630, 0.05% TX-100, 5% normal Goat serum, and 1× PBS).

Primary antibodies were diluted in BB and incubated with samples according to the antibody used. For outer-mitochondria/outer-mitochondria 2-color samples, mouse anti-GFP (Invitrogen, Cat# A11120, used at 1:500) was incubated on samples at 4 °C overnight followed by rabbit anti-TOM20 (Abcam, Cat# ab78547, used at 1:1000) on samples at room temperature for 1 hour. For outer-mitochondria/nucleoid two-color samples, mouse anti-dsDNA (Abcam, Cat# ab27156, used at 1:1000) was incubated with samples at 4 °C overnight followed by rabbit anti-TOM20 (used at 1:500) on samples at room temperature for 1 hour the following day. When labeling inner and outer mitochondria, the inner mitochondria primary and secondary labeling was completed before beginning labeling the outer mitochondria. After each antibody labeling, samples were washed in **WB** (0.05% CA-630, 0.05% TX-100, 0.2% BSA, and 1× PBS) three times for 5 minutes. Two-color samples were labeled with AF647 and CF660C secondary antibodies. Post-fixation was performed using 3% PFA + 0.1% GA in PBS for 10 minutes. Samples were rinsed three times in PBS before being stored in PBS at 4 °C.

**Synaptonemal complex labeling.** After being fixed with 4% PFA, the samples were then washed with 1× PBS 3 times followed by a permeabilization step using 0.5% TX-100 in 1× PBS for 10 minutes at room temperature. Samples were then rinsed in 0.1% TX-100 in PBS and treated with Image-iT Signal Enhancer (Molecular Probes, Cat# I36933) for 30 minutes at room temperature. After 3 washes in 0.1% TX-100 in PBS, the samples were incubated in blocking buffer (0.05% TX-100, 5% normal goat serum, 0.05%, in 1× PBS) for 30 minutes at room temperature. For SYCP3 and SYCP1-C (or SYCP1-N) two-color samples, rabbit anti-SYCP1-C (used at 1:500) (or rabbit anti-SYCP1-N used at 1:500) was incubated with samples at 4 °C overnight followed by mouse anti-SYCP3 (used at 1:500) on samples at room temperature for 1 hour the following day. After each primary antibody labeling, samples were washed in **WB** (0.05% CA-630, 0.05% TX-100, 0.2% BSA, and 1× PBS) three times for 5 minutes. Two-color samples were labeled with AF647 and CF660C secondary antibodies. For SYCP3/SYCP1-C/LaminB three-color samples, rabbit anti-SYCP1-C (used at 1:500) was incubated with samples at 4 °C overnight. The next day mouse anti-SYCP3 (used at 1:500) was incubated with samples at room temperature for 1 hour followed by a 1-hour incubation at room temperature with chicken anti-LaminB (used at 1:200). After each primary antibody labeling, samples were washed in **WB** (0.05% CA-630, 0.05% TX-100, 0.2% BSA, and 1× PBS) three times for 5 minutes. Three-color samples were labeled with CF680 (used at 1:1000), DY650 (used at 1:1000), and Dyomics 634 (used at 1:200) secondary antibodies.

**Golgi labeling**. Since we did not identify a good anti-ManII antibody suitable for immunolabeling, we electroporated HeLa cells with a ManII-GFP plasmid to image the *medial* Golgi apparatus. Cells were transferred to cleaned coverglass and expressed the plasmid for 18-24 hours before being fixed with 4% PFA in 1× PBS for 15 minutes at room temperature. Cells were permeabilized with **PB** (0.3% CA-630, 0.05% TX-100, 0.1% BSA, and 1× PBS) for 3 minutes at room temperature, rinsed three times in PBS, and then blocked in **BB** (0.05% CA-630, 0.05% TX-100, 5% normal Goat serum, and 1× PBS).

Primary antibodies were diluted in BB and incubated with samples according to the antibodies used. For *cis/medial/trans* labeled samples, mouse anti-p230 (BD Bioscience, Cat# 611280, used at 1:1000) and nanobody GFP-binding protein, conjugated with DL650, were incubated with samples at 4 °C overnight followed by rabbit anti-GRASP65 (Abcam, Cat# ab174834, used at 1:2000) on samples for 1 hour at room temperature the next day. Alternatively, for *cis/cis/medial* labeled samples, mouse anti-GM130 (BD Bioscience, Cat# 610822, used at 1:500) and nanobody GFP-binding protein, conjugated with DL650, were incubated on samples together at 4 °C overnight followed by rabbit anti-GRASP65 (used at 1:3000) on samples for 1 hour at room temperature the next day. After each antibody labeling, samples were washed in **WB** (0.05% CA-630, 0.05% TX-100, 0.2% BSA, and 1× PBS) three times for 5 minutes. After labeling with both primary antibodies and nanobody, samples were incubated with secondary antibodies labeled with DY634 and CF680 together. Samples were post-fixed using 3% PFA + 0.1% GA in PBS for 10 minutes. Samples were rinsed three times in PBS before being stored in PBS at 4 °C.

**ER-PM contact sites labeling.** COS-7 cells were grown on Poly-L-Lysine coated glass and transfected with GFP-Sec61β, GFP-ORP5, mCherry-ORP5, GFP-ESYT2, or mCherry-ESYT2 (in different combinations) using Lipofectamine2000. Cells were fixed with 3% PFA + 0.1% GA 18-24 hours after transfection. If the plasma membrane was lectin-labeled, the samples were labeled directly after fixation but before permeabilization. Cells were rinsed with Hanks balanced salt solution (HBSS; Gibco, Cat# 14025-092) three times before labeling with 1 μg/mL WGA-CF680 (Biotium, 29029-1) diluted in HBSS for 10-30 minutes at room temperature. Cells were then rinsed three times in HBSS and once in PBS before being permeabilized using **PB** (0.3% CA-630, 0.05% TX-100, 0.1% BSA, and 1× PBS) for 3 minutes at room temperature. Samples were rinsed three times in PBS and blocked for at least 1 hour with **BB** (0.05% CA-630, 0.05% TX-100, 5% normal Goat serum, and 1× PBS). Primary antibodies and/or nanobodies in BB were incubated with samples overnight at 4 °C.

For two-color samples labeling ER and PM (expressing GFP-Sec61β) and two-color samples labeling contact proteins and PM (expressing GFP-ORP5 or GFP-ESYT2), GFP was labeled with rabbit anti-GFP (used at 1:500 overnight at 4 °C) followed by a secondary AF647 antibody. Some ER and PM samples also expressed mCherry-ORP5 or mCherry-ESYT2, which was not immunolabeled. For two-color samples labeling ER and contact proteins (expressing GFP-Sec61β with either mCherry-ORP5 or mCherry-ESYT2), rabbit anti-mCherry (Abcam, Cat# ab167453 used at 1:500 or 1:1000 overnight at 4 °C) and mouse anti-GFP (used at 1:500 overnight 4 °C) were used to label the fluorescent proteins, followed by secondary antibodies labeled with AF647 and CF660C. For three-color samples labeling ER, contact site proteins, and PM (expressing GFP-Sec61β with either mCherry-ORP5 or mCherry-ESYT2), WGA was used to label the PM, rabbit anti-mCherry (used at 1:500 or 1:1000 overnight at 4 °C) and mouse anti-GFP (used at 1:500 overnight 4 °C) were used to label the fluorescent proteins, followed by secondary antibodies labeled with DY634 and DL650. Post-fixation was performed using 3% PFA + 0.1% GA in PBS for 10 minutes. Samples were rinsed three times in PBS before being stored in PBS at 4 °C.

**Mitochondria and microtubule**. COS-7 cells were used for two-color labeling of mitochondria and microtubules. Cells were grown on cleaned and Poly-L-Lysine coated glass before being fixed in 3% PFA + 0.1% GA in 1× PBS for 15 minutes. Samples were rinsed three times in PBS before permeabilizing for 3 minutes using permeabilization buffer (**PB**; 0.3% CA-630, 0.05% Triton X-100, 0.1% BSA, and 1× PBS) at room temperature. Samples were then rinsed three more times with PBS followed by 1 hour in block buffer (**BB**; 0.05% CA-630,0.05% Triton X-100, 5% normal Goat serum, and 1× PBS). Primary antibodies rabbit anti-TOM20 (used at 1:500) and mouse anti-tubulin (used at 1:1000) were diluted in BB and incubated with samples overnight at 4 °C. Samples were then washed in wash buffer (**WB**; 0.05%

CA-630,0.05% TX-100, 0.2% BSA, and 1× PBS,) three times for 5 minutes each before secondary antibody labeling for 1 hour at room temperature diluted in BB. Samples were then washed in WB three times for 5 minutes each. Post-fixation was performed using 3% PFA + 0.1% GA in PBS for 10 minutes. Samples were rinsed three times in PBS before being stored in PBS at 4 °C.

**Imaging buffer and sample mounting.** The conventional β-mercaptoethanol (βME) STORM imaging buffer was prepared as previously reported (*24*). The imaging buffer was made every time immediately before use where catalase and glucose oxidase were diluted in base buffer (50 mM Tris pH 8.0, 50 mM NaCl, 10% glucose) with the addition of βME (Sigma-Aldrich, Cat# M3148-25ML). The final concentration of βME is 143 mM. The samples were mounted in a custom-designed sample holder as previously described (*8*). Briefly, the sample coverslip was mounted in the sample holder facing up. Then 100 μL of imaging buffer was evenly spread on the sample coverslip and a clean coverslip was put on top (attention was given to avoid bubbles trapped between the two coverslips). Excess imaging buffer was drained using Kimwipes. The samples were then sealed with two-component silicone glue (Picodent Twinsil, Picodent, Wipperfürth, Germany). After the silicone glue hardened (typically 20-30 min), the samples were transferred to the 4Pi-SMS microscope for imaging. The imaging buffer would usually allow for imaging of about 8-10 hours.

#### Multi-color 4Pi-SMS setup

The multi-color 4Pi-SMS system was built based on the previously described instrument (*8*) with minor modifications (fig. S1). The oil immersion objectives were replaced with silicone immersion objectives (100 x/1.35, Olympus) for better refractive index matching. The system was equipped with an excitation laser at 642 nm (MPB Communications, 2RU-VFL-2000-642-B1R) and an activation laser at 405 nm (Coherent OBIS 405 LX, 50 mW). Details about the dichroic beamsplitter and emission filters used in the system are shown in fig. S1B. The conventional fluorescence follows the same emission path as the previous design and is collected by a sCMOS camera (ORCA-Flash 4.0v2, Hamamatsu) (Camera 1, fig. S1A). The salvaged fluorescence is reflected by the dichroic beamsplitter to the back side of the system and collected by an EMCCD camera (128 × 128 pixels; iXon DU860, Andor) (Camera 2, fig. S1C). Both cameras were controlled by custom-written LabVIEW (National Instruments) programs.

#### Image acquisition

During image acquisition, the sCMOS camera was set to external trigger mode and the EMCCD camera was set to internal trigger and frame transfer mode. For synchronization, the Fire output of the EMCCD camera was used to trigger the sCMOS camera. The electron multiplication gain of the EMCCD camera was set to 200 for all experiments. Biological samples were imaged at 100 Hz with a laser (642 nm) intensity of about 7.5 kW/cm$^2$ (two-color imaging) or 200 Hz at about 15 kW/cm$^2$ (three-color imaging). The 405-nm activation laser was manually adjusted to maintain a low density of single molecules per frame. For samples thicker than 1 μm, the stage was translated at 500-nm steps every 3000 frames for multiple times to cover the entire volume. Typically, 180,000 to 360,000 frames were recorded which corresponds to acquisition times of 15 min to 1 hr.

#### Image analysis and color assignment

The images recorded with the sCMOS camera (conventional fluorescence) were analyzed as previously described (*8*). Briefly, the lateral positions (xy) of single molecules were determined by fitting with sCMOS-specific algorithms (*24*). The phase values (z positions) were estimated from the $0^{th}$ moment Gaussian intensities of the four images and unwrapped using the metric developed previously (*8*). To translate the phase values to axial positions, the intensity modulation frequency for each dye was determined from the simulated 4Pi PSFs using a pupil function-based approach (*25*). The simulation was performed using the average wavelength of the detected emission spectra in the conventional fluorescence channel of each dye (fig. S3, C to G). For the images recorded with the EMCCD camera (salvaged fluorescence), the readout counts were converted to photoelectrons using the conversion factor provided by the manufacturer. Then the pixel-wise median operation (time window = 3000 frames) in MATLAB (MathWorks) was used as a filter algorithm to remove the background signals. The positions of localized molecules in the sCMOS images were then mapped to the EMCCD images using a second-order affine transformation matrix obtained by imaging a fluorescent bead sample. The corresponding regions in the EMCCD images were cropped out (5 × 5 pixels, 170 nm pixel size) and multiplied by a normalized 2D Gaussian distribution with a standard deviation of 130 nm centered at the positions of the molecules. This Gaussian weighting was performed to reduce the influence from the background regions. The intensity of the molecules in the EMCCD images was calculated by summing the values of the Gaussian-weighted regions. For color assignment, the intensities of the molecules in the salvaged and conventional fluorescence images were plotted on a logarithmic scale (Fig. 1, D and H) and binned to a 2D histogram intensity image (fig. S4G and fig. S5J). An appropriate threshold (typically 2% of each peak value) was applied to separate the different dye molecules with a low cross-talk (fig. S4, G and H; fig. S5, J and K). After color assignment, the phase-unwrapped values of each dye molecules from the sCMOS images were translated to axial positions using the wavelength-dependent modulation frequencies obtained above. Localizations (x, y, and z) from all color channels were combined for drift correction using a custom algorithm based on redundant cross-correlation (*26-28*). For samples requiring axial stepping, multiple optical sections were aligned using a 3D cross-correlation method (*26*). Localizations that appear at consecutive frames within a radius of 2 times the localization precision were considered to represent the same molecule and combined. All 4Pi-SMS images and videos were rendered with Vutara SRX software (Bruker).

#### Thickness and separation measurements of the Golgi cisternae

For each 3D Golgi volume, we cropped 1-µm wide subregions along the x- or y-direction. We manually reoriented the subregions to generate 2D projection images providing a side view of the region of the Golgi stack to be analyzed (Fig. 3, F and G; fig. S10, E and F). Next, we took line profiles (thickness = 100 nm) at positions where the three markers are parallel to each other (Fig. 3F and fig. S10F). Each intensity profile was fit by a Gaussian function to determine the thickness (estimated as the full-width at half-maximum of the Gaussian distribution) and the separation between cisternae (distance between peak positions). The average separation between GRASP65 and GM130 is not significantly different from zero (student's $t$-test, $p = 0.6$).

#### Quantification of the ER-PM contact sites images

The two-color images of the contact site proteins (ORP5 or ESYT2) and PM (fig. S11, B and C) were automatically divided into 100 × 100 nm x-y subregions. To ensure a high signal-to-noise ratio, only subregions with more than 100 localizations from the PM labeling (WGA) and 300 localizations from ORP5 (or ESYT2) were kept for further data analysis (more than 2800 subregions for each condition). The WGA labeling featured much brighter staining of the top PM compared to the bottom PM which resulted in much higher WGA localization densities at the top PM than the bottom PM. We therefore used the top PM signal only for distance measurements. WGA localizations with z positions below the median z positions of ORP5 (or ESYT2) were rejected as they were considered to belong to the bottom PM. The remaining z positions of WGA and the z positions of ORP5 (or ESYT2) were binned into histograms, respectively. The histograms of WGA and ORP5 (or ESYT2) localizations were both fit by Gaussian functions to determine the two peak positions and the distance between them (Fig. 4G).

#### Simulation of the SF approach performance

The salvaged fluorescence (SF) signal is mainly determined by the transition wavelength of the dichroic beamsplitter while the conventional fluorescence signal is determined by the spectral window confined by the dichroic beamsplitter and the emission filter (in front of Camera 1; fig. S1A). The performance of the SF approach is therefore expected to depend on the choice of the transition wavelength of the dichroic beamsplitter. To investigate this phenomenon, we performed simulations for transition wavelengths ranging from 650 nm to 680 nm (fig. S2). To use realistic simulation parameters, we based our simulation on experimental data. For each dye, we randomly selected 2 million single molecules from real imaging experiments. The photon number of each molecule (determined from the conventional fluorescence channel) was converted to the total photon number representing the whole emission spectrum taking the experimentally used spectral detection window into account. Next, we took the transition profile of a commercially available dichroic beamsplitter (Semrock Di01-R405/488/561/635) which features a steep rising edge in the ~657 nm range. The transition wavelength of the dichroic beamsplitter was defined as the wavelength at which the transmission is 50%. The transmission profile starts to rise at 650 nm. Below that value, we assumed 2% transmission. Above 664 nm, we assumed 98% transmission. For our simulation, we shifted this transition profile across the spectrum starting at a transition wavelength of 650 nm and ending at 680 nm, with the assumption that the conventional fluorescence channel collects the transmitted signal up to 750 nm and the salvaged fluorescence channel collects the reflected signal upwards from 645 nm (based on the used excitation laser wavelength of 642 nm). The fraction of conventional and salvaged fluorescence was calculated for each simulation and photon numbers were allocated to each channel accordingly. We then generated synthetic single-molecule images based on these photon numbers in each channel and added background and readout noise similar to experimentally observed values. Finally, these images were analyzed in the same way as experimental data and the cross-talk and localization precisions were calculated taking advantage of the ground truth provided by the simulation and averaged for 2 million simulated molecules each.

We found that less than 10% of Alexa 647 (when compared to CF660C) and DL650 (when compared to CF680) molecules were rejected at transition wavelengths of 661 nm or above, and 662 nm or above, respectively. As expected, localization precision degraded with increasing transition wavelengths since larger transition wavelengths lead to narrower conventional fluorescence spectral windows. Up to 670 nm transition wavelength, the loss in localization precision stayed below 2 nm, which we considered acceptable. The dichroic beamsplitter used in our microscope (Chroma ZT405/488/561/647rpc) features a transition wavelength of 668 nm which falls within this acceptable window between 661 nm or 662 nm on the lower end and 670 nm on the upper end.

Although we selected commercially available dichroic beamsplitters and emission filters in this work, it is possible to increase the collection efficiency of the conventional fluorescence channel by using custom dichroic beamsplitters with a transition edge closer to the excitation wavelength. For two-objective systems, collecting salvaged fluorescence at both objectives can further improve the performance. Additionally, the detected salvaged fluorescence can be used to improve the localization precision but at the cost of image registration error and added complexity in the data analysis.

#### Supplementary Figures

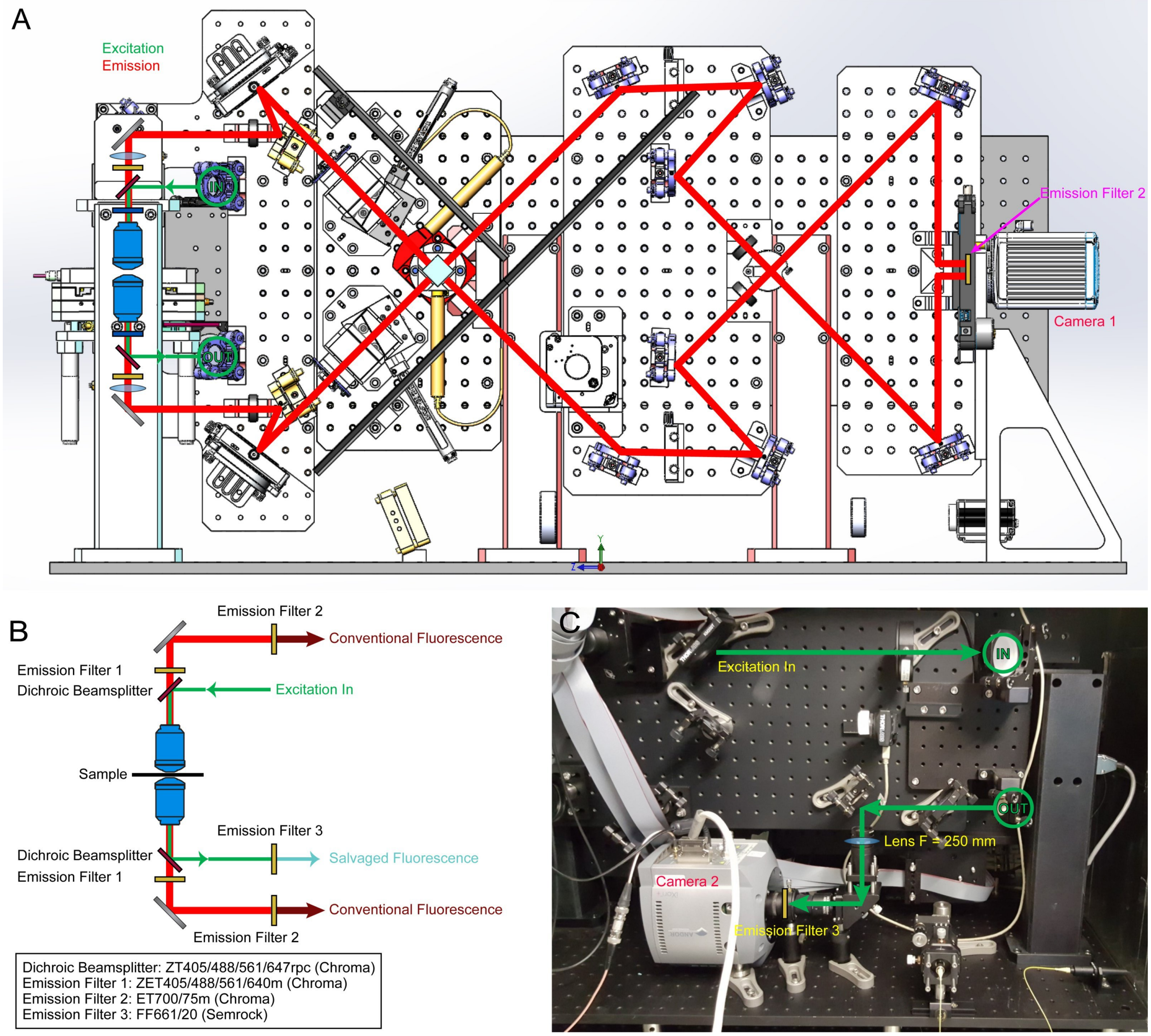


**Fig. S1. Multicolor 4Pi-SMS microscope design and setup**. (A) Schematic drawing of the front of the 4Pi-SMS system. The excitation laser (green solid line) emerges from the back side of the system (IN), is reflected by the upper dichroic beamsplitter, passes through the objectives and sample, is reflected by the lower dichroic beamsplitter and returns to the back side of the system (OUT). The conventional fluorescence (red solid line) is collected by camera 1. (B) Schematic of positions and information about the dichroic beamsplitters and emission filters used in the system. (C) Photograph of the backside of the 4Pi-SMS system. The positions where the excitation laser goes to (IN) and comes back from (OUT) the front of the system are marked. The salvaged fluorescence is collected by camera 2.

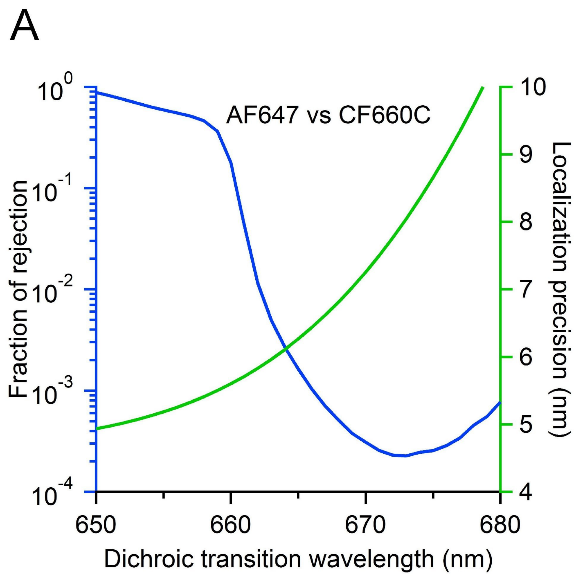


**Fig. S2. Simulation of the SF approach performance for different transition wavelengths of the dichroic beamsplitter.** (A) Simulation results for AF647 and CF660C, the dyes used in two-color imaging. The blue line shows the fraction of rejected molecules when 1% cross-talk is achieved. The green line shows the localization precision of AF647. (B) Simulation results for DL650 and CF680, two of the dyes used in three-color imaging. The blue line shows the fraction of rejected molecules when 2% cross-talk is achieved. The green line shows the localization precision of DL650.

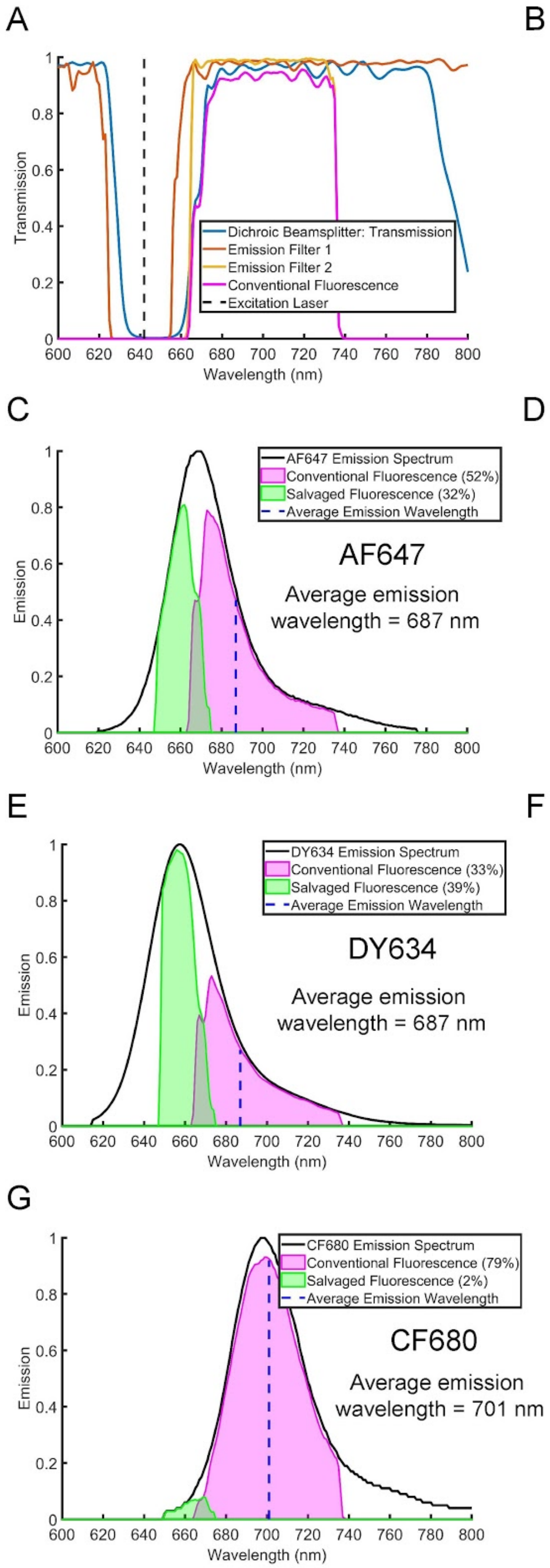
A
1
0.8
0.6
0.4
0.2
0
Transmission
Dichroic Beamsplitter: Transmission
Emission Filter 1
Emission Filter 2
Conventional Fluorescence
Excitation Laser
600 620 640 660 680 700 720 740 760 780 800
Wavelength (nm)
B
Dichroic Beamsplitter: Reflection
Emission Filter 3
Salvaged Fluorescence
Excitation Laser
C
Emission
AF647 Emission Spectrum
Conventional Fluorescence (52%)
Salvaged Fluorescence (32%)
Average Emission Wavelength
AF647
Average emission
wavelength = 687 nm
D
CF660C Emission Spectrum
Conventional Fluorescence (75%)
Salvaged Fluorescence (9%)
Average Emission Wavelength
CF660C
Average emission
wavelength = 693 nm
E
DY634 Emission Spectrum
Conventional Fluorescence (33%)
Salvaged Fluorescence (39%)
Average Emission Wavelength
DY634
Average emission
wavelength = 687 nm
F
DL650 Emission Spectrum
Conventional Fluorescence (67%)
Salvaged Fluorescence (17%)
Average Emission Wavelength
DL650
Average emission
wavelength = 692 nm
G
CF680 Emission Spectrum
Conventional Fluorescence (79%)
Salvaged Fluorescence (2%)
Average Emission Wavelength
CF680
Average emission
wavelength = 701 nm

**Fig. S3. Transmission profiles for the conventional and salvaged fluorescence.** (A) Transmission profiles of the dichroic beamsplitter and emission filters for the conventional fluorescence beam path. The magenta line shows the combined transmission profile of the dichroic beamsplitter and emission filters. (B) Transmission profiles of the dichroic beamsplitter and emission filters for the salvaged fluorescence beam path. The green line shows the combined profiles. (C to G) The emission spectra and the fraction of conventional and salvaged fluorescence for each dye used in this work. The blue dashed line indicates the average wavelength of each dye in the detected spectral window of the conventional fluorescence channel. The numerical value is listed in each panel.

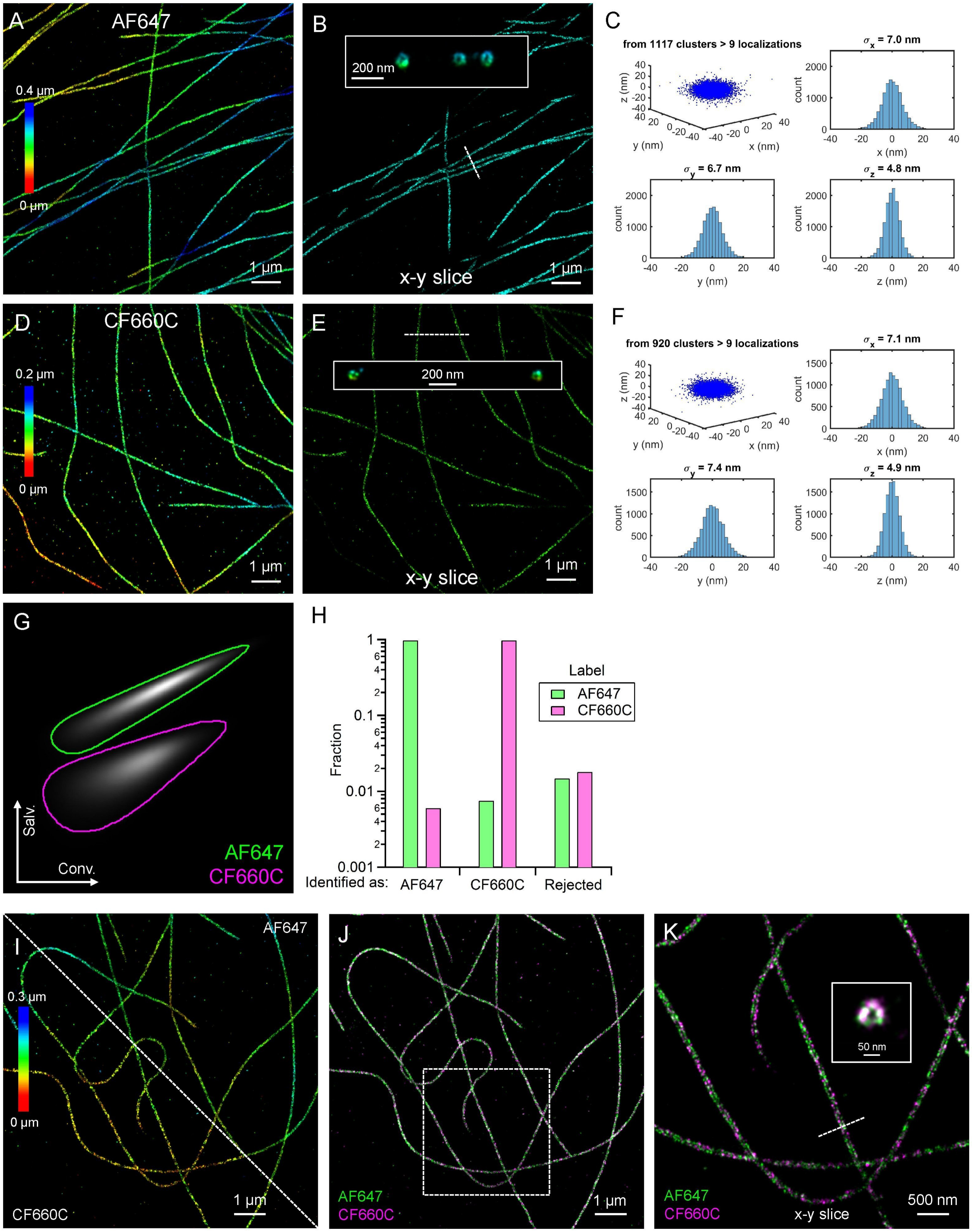


**Fig. S4. Two-color 4Pi-SMS imaging performance on microtubule samples.** (A to F) 4Pi-SMS images of microtubules labeled with AF647 or CF660C in COS-7 cells and the 3D localization precision. (A, D) z-projections of the data sets. (B, E) 20-nm thick x-y slices of the same data. (C, F) Clusters of localizations from the same dye molecule (emitters that appeared in consecutive frames within a radius of two times the localization precision) were aligned by their center of mass to generate the shown 3D plots. Histograms of the distributions in x, y, z were fit with Gaussian functions, and the standard deviations ($\sigma_x$, $\sigma_y$ and $\sigma_z$) are reported. (G) The binned 2D intensity histogram of AF647 and CF660C from more than 2 million localizations of each dye based on the same data as shown in (Fig. 1D). The plot shows salvaged fluorescence intensity (y-axis) versus conventional fluorescence intensity (x-axis). The solid lines show the threshold where the intensity value is 2% of each peak intensity. (H) Cross-talk between AF647 and CF660C on a logarithmic scale representing the same data as shown in (Fig. 1E). Localizations outside the two threshold boundaries in (G) are rejected. (I) Two-color images of microtubules (anti-α-tubulin antibody labeled with AF647 and CF660C together) in a COS-7 cell. The lower left and upper right corners show the CF660C and AF647 labeling, respectively. (J) Merged image of the two labels in (I). (K) A 20-nm thick x-y slice of the boxed region in (J). Inset: a 100-nm thick cross-section along the dashed line in (K).

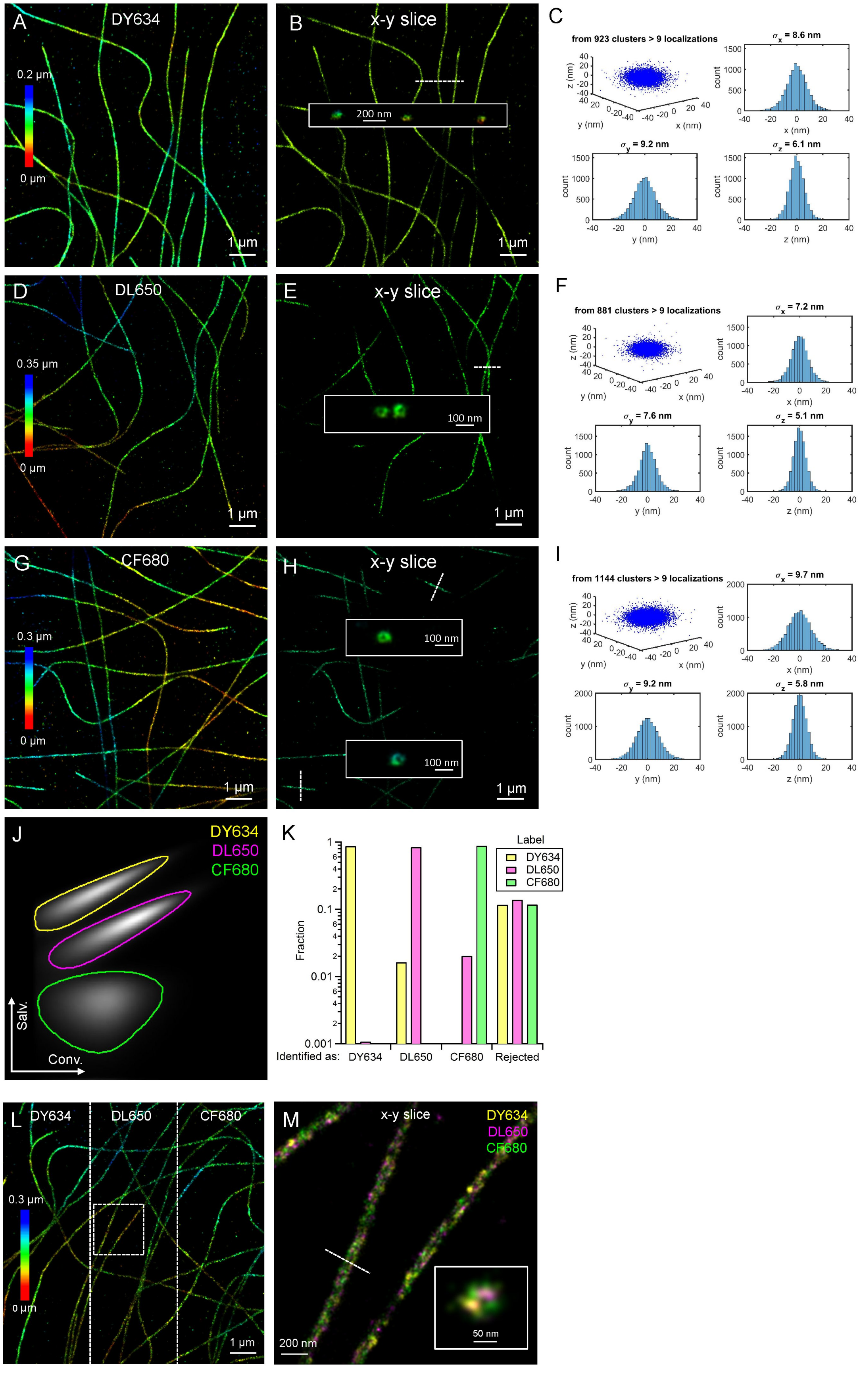
A
DY634
0.2 µm
0 µm
1 µm
B
x-y slice
200 nm
1 µm
C
from 923 clusters > 9 localizations
σx = 8.6 nm
σy = 9.2 nm
σz = 6.1 nm
D
DL650
0.35 µm
0 µm
1 µm
E
x-y slice
100 nm
1 µm
F
from 881 clusters > 9 localizations
σx = 7.2 nm
σy = 7.6 nm
σz = 5.1 nm
G
CF680
0.3 µm
0 µm
1 µm
H
x-y slice
100 nm
100 nm
1 µm
I
from 1144 clusters > 9 localizations
σx = 9.7 nm
σy = 9.2 nm
σz = 5.8 nm
J
DY634
DL650
CF680
Salv.
Conv.
K
Label
DY634
DL650
CF680
Fraction
Identified as: DY634 DL650 CF680 Rejected
L
DY634
DL650
CF680
0.3 µm
0 µm
1 µm
M
x-y slice
DY634
DL650
CF680
200 nm
50 nm

**Fig. S5. Three-color 4Pi-SMS imaging performance on microtubule samples.** (A-I) 4Pi-SMS images of microtubules labeled with DY634, DL650 or CF680 in COS-7 cells and the 3D localization precision. (A, D, G) z-projections of the data sets. (B, E, H) 20-nm thick x-y slices of the same data. (C, F, I) Clusters of localizations from the same dye molecule (emitters that appeared in consecutive frames within a radius of two times the localization precision) were aligned by their center of mass to generate the shown 3D plots. Histograms of the distributions in x, y, z were fit with Gaussian functions, and the standard deviations ($\sigma_x$, $\sigma_y$ and $\sigma_z$) are reported. (J) The binned 2D intensity histogram from more than 2 million localizations of each dye based on the same data as shown in (Fig. 1H). The plot shows salvaged fluorescence intensity (y-axis) versus conventional fluorescence intensity (x-axis). The solid lines show the threshold where the intensity value is 2% of each peak intensity. (K) Cross-talk between each dye pair on a logarithmic scale representing the same data as shown in (Fig. 1I). Localizations outside the three threshold boundaries in (J) are rejected. (L) Three-color image of microtubules co-labeled with all three dyes in a COS-7 cell. (M) Magnified image of the boxed region in (L) with the three color channels overlaid. Inset: a 20-nm thick cross-section taken at the dashed line in (M).

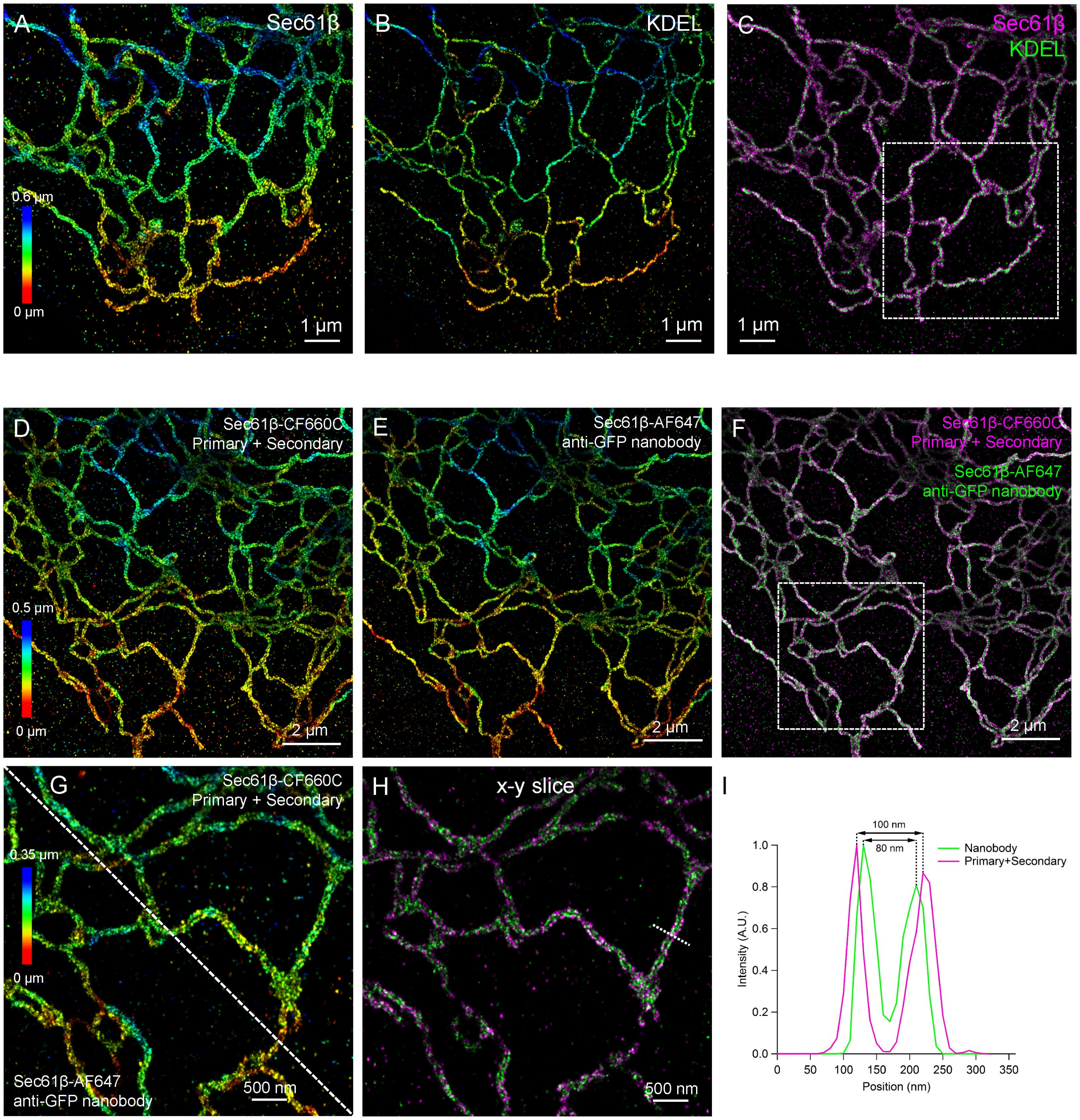


**Fig. S6. Two-color 4Pi-SMS images of ER in COS-7 cells.** (A to C) Two-color images of ER membrane (overexpressed GFP-Sec61β labeled with anti-GFP antibody) and ER lumen (overexpressed mCherry-KDEL labeled with anti-RFP nanobody) (Movie S2). Merged image of the two labels in (A) and (B) is shown in (C). The boxed region in (C) is shown in (Fig. 2B). (D to F) Two-color images of ER membrane (overexpressed GFP-Sec61β labeled both with antibodies and nanobody). (G) Magnified image of the boxed region in (F). The lower left corner shows the nanobody labeling while the upper right corner shows the antibody labeling. (H) A 50-nm thick x-y slice of (F). (I) Intensity profile along the dashed line in (H) averaged over a thickness of 100 nm.

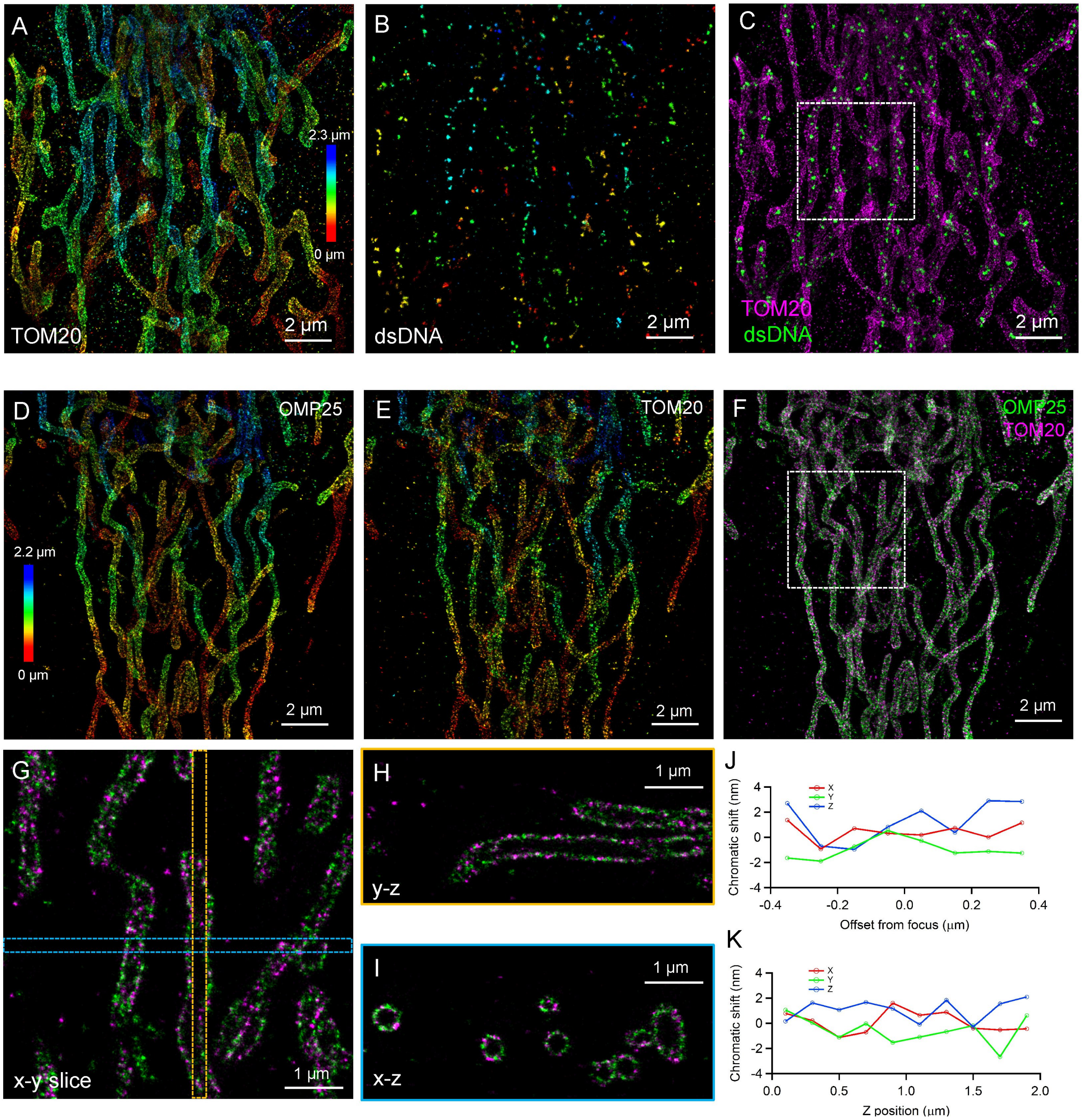


**Fig. S7. Two-color 4Pi-SMS images of mitochondria in HeLa cells.** (A to C) Two-color images of the outer membrane of mitochondria (anti-TOM20 antibody) and mitochondrial DNA (anti-dsDNA antibody) (Movie S3). Merged image of the two labels in (A) and (B) is shown in (C). The boxed region in (C) is shown in (Fig. 2F). (D to F) Two-color image of the outer membrane of mitochondria (overexpressed GFP-OMP25 labeled with anti-GFP antibody alongside anti-TOM20 antibody). The dataset was reconstructed from 4 optical sections with 500-nm step sizes. (G) A 200-nm thick x-y slice of the boxed region in (F). (H) y-z view of the orange dashed box (200 nm wide) in (G). (I) x-z view of the blue dashed box (200 nm wide) in (G). (J) 3D chromatic shifts at different offset from the focus center (100-nm thick volumes) of each optical section averaged over the 4 sections. (K) 3D chromatic shifts at different z-positions from non-overlapping 200-nm thick volumes after the 4 optical sections were aligned and combined. Chromatic shifts stay within a 3-nm range in both cases.

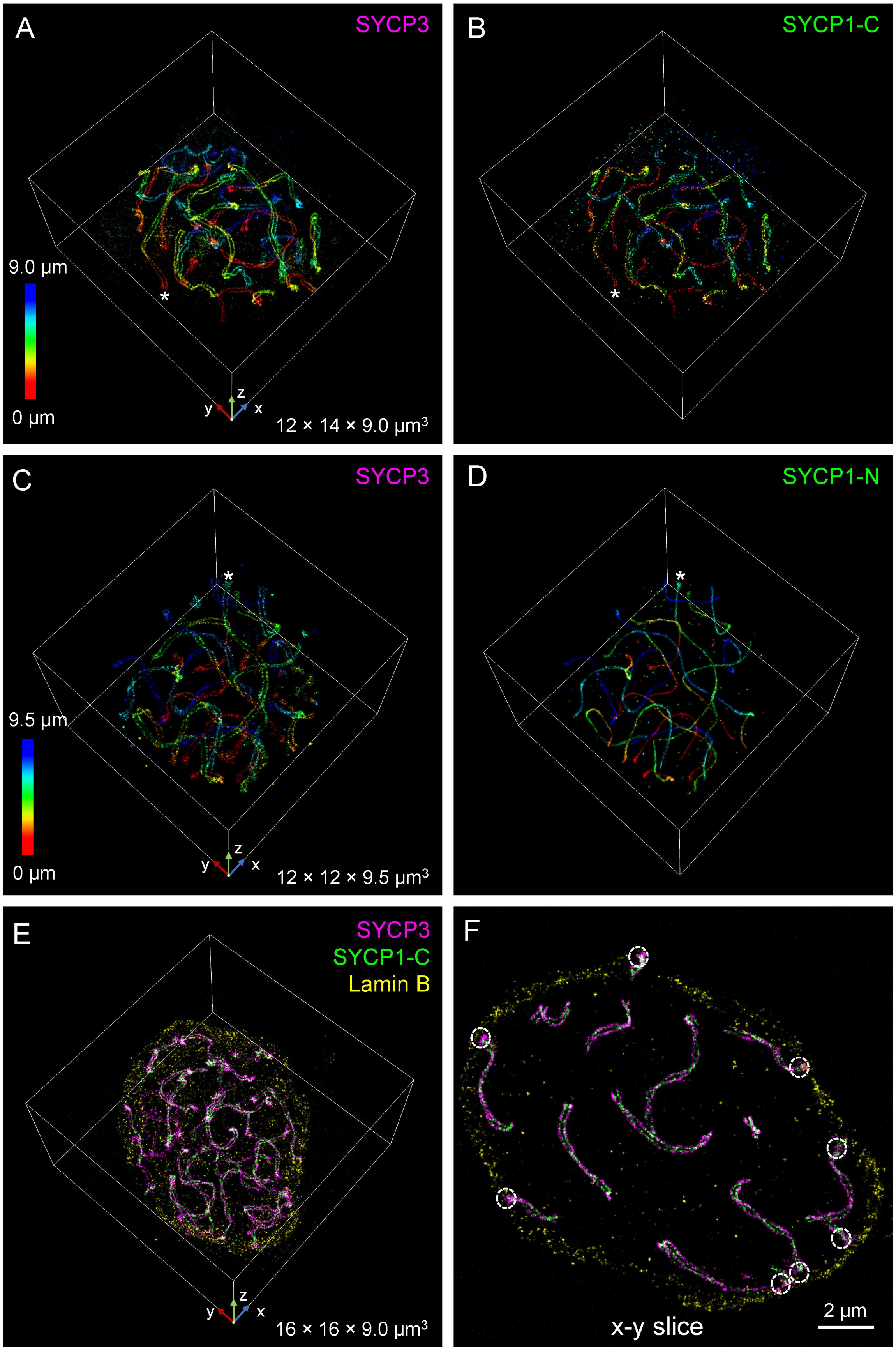
A
SYCP3
9.0 µm
0 µm
z
y
x
12 × 14 × 9.0 µm3
B
SYCP1-C
C
SYCP3
9.5 µm
0 µm
z
y
x
12 × 12 × 9.5 µm3
D
SYCP1-N
E
SYCP3
SYCP1-C
Lamin B
z
y
x
16 × 16 × 9.0 µm3
F
x-y slice
2 µm

**Fig. S8. Multicolor 4Pi-SMS images of synaptonemal complexes in mouse spermatocytes.** (A and B) Two-color images of synaptonemal complexes labeled with anti-SYCP3 antibody and anti-SYCP1 C-terminal antibody (Movie S4, part I). 3D overlay of the synaptonemal complex marked by a white asterisk in (A) and (B) is shown in (Fig. 2J). (C and D) Two-color images of synaptonemal complexes labeled with anti-SYCP3 antibody and anti-SYCP1 N-terminal antibody (Movie S4, part II). 3D overlay image of the synaptonemal complex marked by a white asterisk in (A) and (B) is shown in (Fig. 2M). (E) Three-color image of synaptonemal complexes labeled with anti-SYCP3 antibody, anti-SYCP1 C-terminal antibody and anti-Lamin B antibody (Movie S4, part III). (F) A 2-μm thick x-y slice through the middle of the volume in (E). The white dashed circles indicate the positions where synaptonemal complexes connect to the nuclear lamina. Synaptonemal complexes not marked by dashed circles make contact with the nuclear lamina above or below the slice. The size of the white 3D boundary boxes (x × y × z) is provided in the lower right corner of the images (A, C, E).

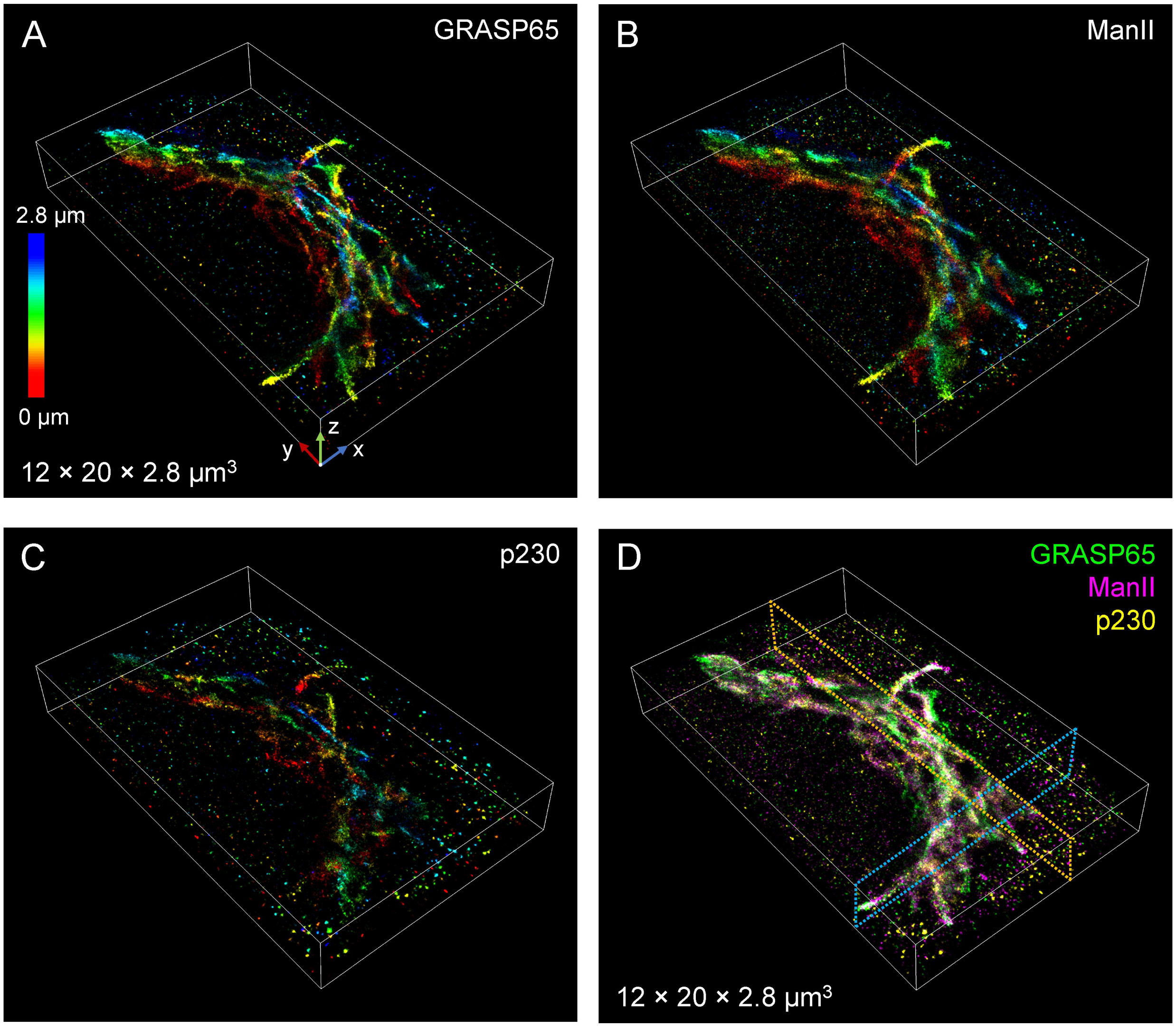


**Fig. S9. Three-color 4Pi-SMS images of the Golgi apparatus using *cis*, *medial* and *trans* markers in a HeLa cell.** (A to D) 3D images of *cis* Golgi (anti-GRASP65 antibody), *medial* Golgi (overexpressed ManII-GFP and labeled with anti-GFP nanobody), and *trans* Golgi (anti-p230 antibody) markers (Movie S5, part I). Merged image of the three labels in (A to C) is shown in (D) and is the same as (Fig. 3A). The size of the white 3D boundary boxes (x × y × z) is provided in the lower left corner of the images (A and D).

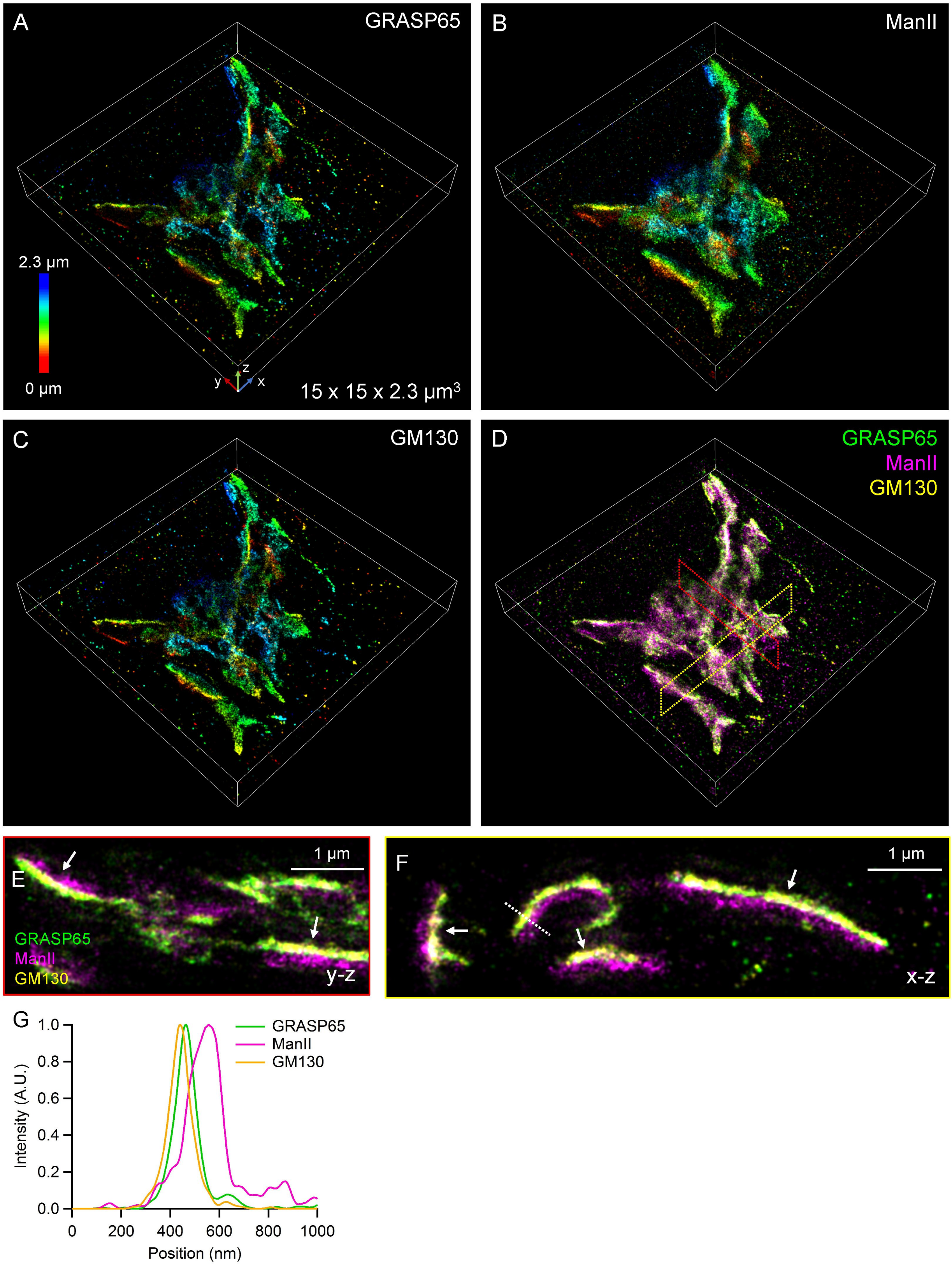

A
GRASP65
2.3 µm
0 µm
15 x 15 x 2.3 µm3
B
ManII
C
GM130
D
GRASP65
ManII
GM130
E
1 µm
GRASP65
ManII
GM130
y-z
F
1 µm
x-z
G
Intensity (A.U.)
Position (nm)
GRASP65
ManII
GM130

**Fig. S10. Three-color 4Pi-SMS images of the Golgi apparatus using one *medial* and two *cis* markers in a HeLa cell.** (A to D) 3D overview images of *cis* Golgi (anti-GRASP65 antibody), *cis* Golgi (anti-GM130 antibody), and *medial* Golgi (overexpressed ManII-GFP labeled with anti-GFP nanobody) (Movie S5, part II). The size of the white 3D boundary boxes (x × y × z) is provided in the lower right corner of the images (A). (E) 1-µm thick y-z cross-section centered at the red dashed region in (D). (F) 1-µm thick x-z cross-section centered at the yellow dashed region in (D). (G) Intensity profile along the dashed line in (F).

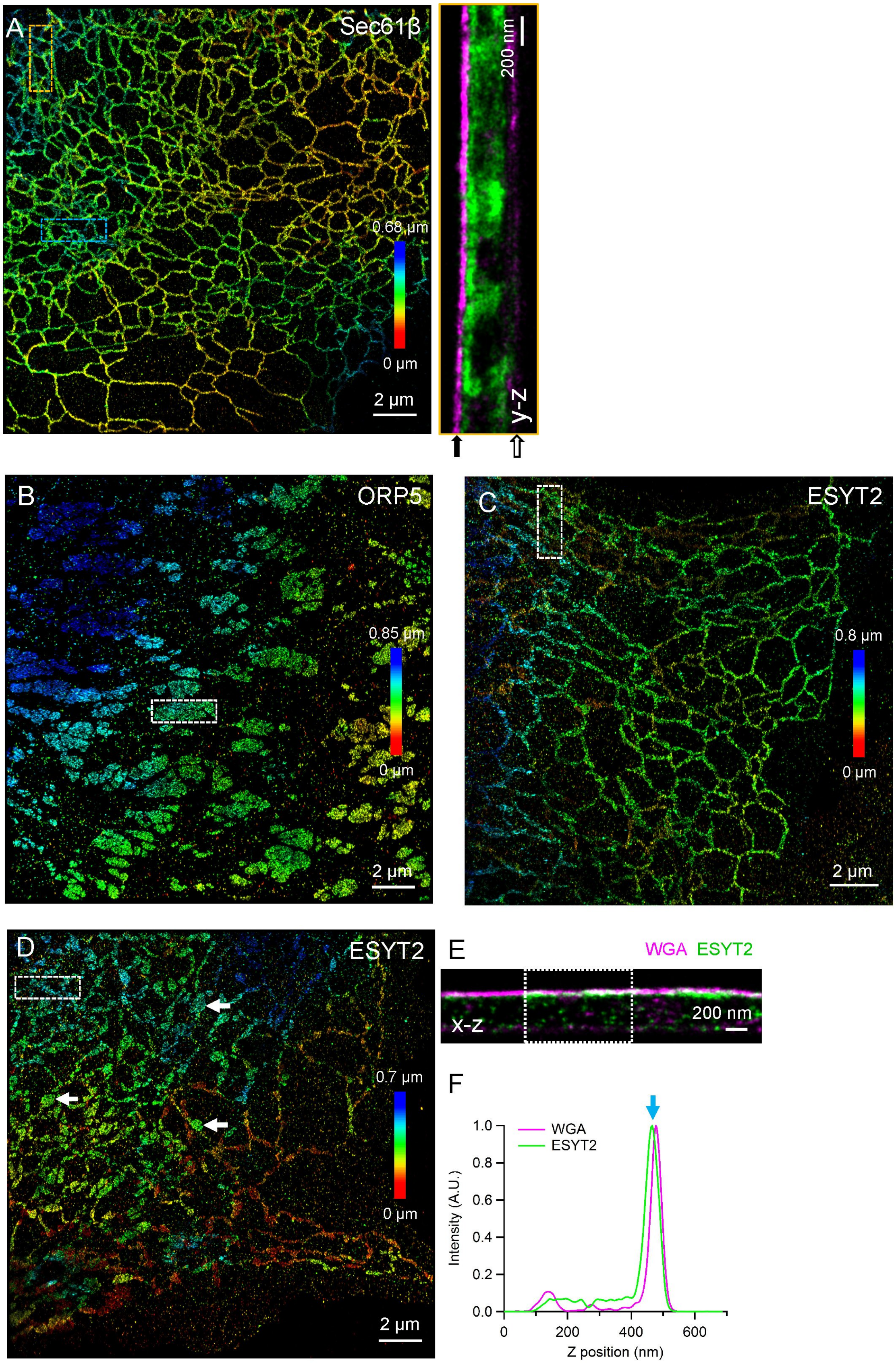

A
Sec61β
0.68 μm
0 μm
2 μm
200 nm
y-z
B
ORP5
0.85 μm
0 μm
2 μm
C
ESYT2
0.8 μm
0 μm
2 μm
D
ESYT2
0.7 μm
0 μm
2 μm
E
WGA ESYT2
x-z
200 nm
F
WGA
ESYT2
Intensity (A.U.)
1.0
0.8
0.6
0.4
0.2
0.0
0
200
400
600
Z position (nm)

**Fig. S11. Two-color 4Pi-SMS images of the ER and contact site proteins in COS-7 cells.** (A) Two-color image of ER (overexpressed GFP-Sec61β labeled with anti-GFP antibody) and PM (labeled with WGA) in a cell overexpressing mCherry-ESYT2 (Movie S6, part III). y-z view is taken from the boxed regions shown in blue and yellow, respectively (3 x 1 μm). (B) Image of ORP5 (overexpressed GFP-ORP5 labeled with anti-GFP antibody) from a two-color image with the PM. x-z view of the boxed region in (B) is shown in (Fig. 4C). (C) Image of ESYT2 (overexpressed GFP-ESYT2 labeled with anti-GFP antibody) from a two-color image with the PM. y-z view of the boxed region in (C) is shown in (Fig. 4E). (D) Two-color image of ESYT2 (overexpressed GFP-ESYT2 labeled with anti-GFP antibody) and the PM (labeled with WGA). The white arrows indicate some small patch-like structures of ESYT2. (E) x-z view of the boxed region (3 x 1 μm) in (D). (F) Axial intensity profile averaged across the dashed box in (E). The blue arrow indicates the small distance between WGA and ESYT2.

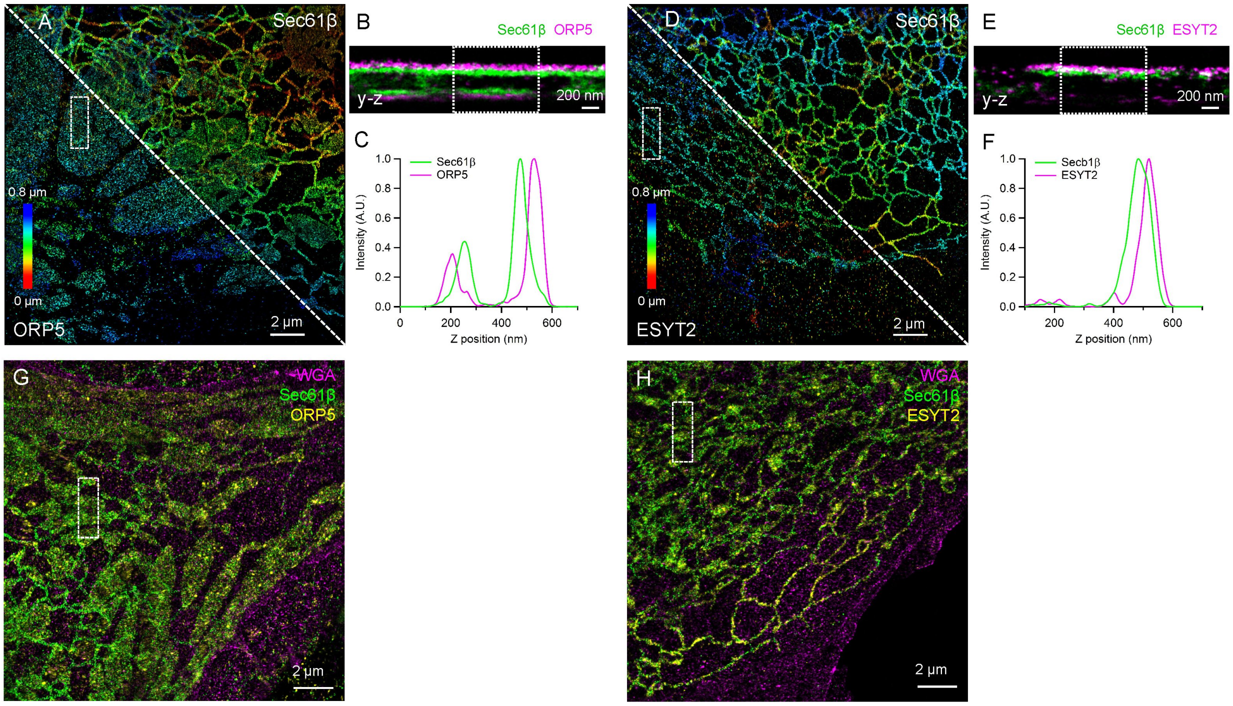


**Fig. S12. Multicolor 4Pi-SMS images of the ER and contact site proteins in COS-7 cells.** (A) Two-color image of the ER membrane (overexpressed GFP-Sec61β labeled with anti-GFP antibody; shown in the upper right half) and ORP5 (overexpressed mCherry-ORP5 labeled with anti-mCherry antibody; shown in the lower left half). (B) y-z view of the boxed region (3 x 1 µm) in (A). (C) Axial intensity profile averaged across the dashed box in (B). (D) Two-color image of the ER membrane (overexpressed GFP-Sec61β labeled with anti-GFP antibody; shown in the upper right half) and ESYT2 (overexpressed mCherry-ESYT2 labeled with anti-mCherry antibody; shown in the lower left half). (E) y-z view of the boxed region (3 x 1 µm) in (D). (F) Axial intensity profile averaged across the dashed box in (E). (G) Three-color image of Sec61β, ORP5 and WGA (Movie S7, part I). y-z view of the boxed region in (G) is shown in (Fig. 4H). (H) Three-color image of Sec61β, ESYT2 and WGA (Movie S7, part II). y-z view of the boxed region in (H) is shown in (Fig. 4J).

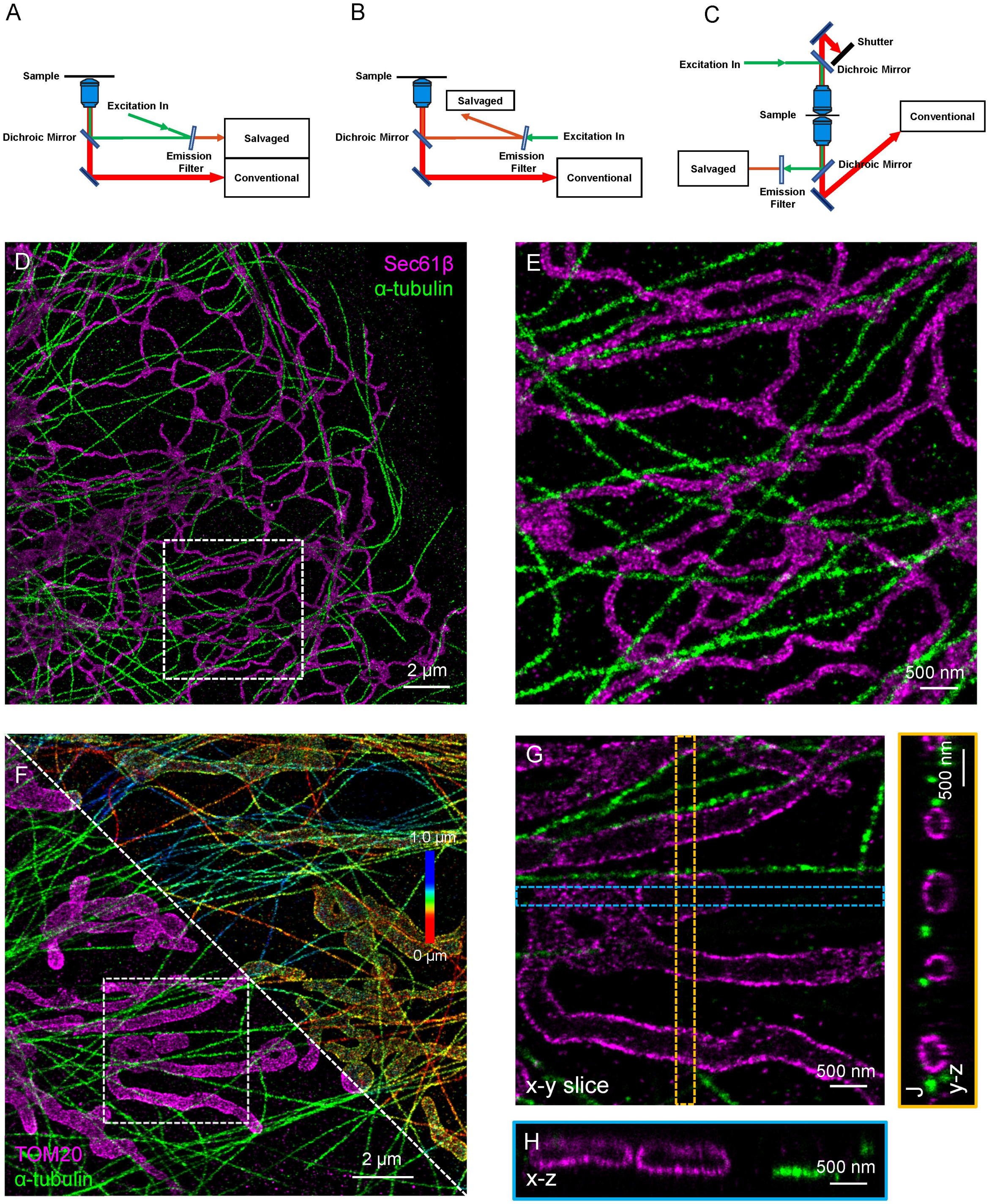


**Fig. S13. Alternative implementations of salvaged fluorescence detection and single-objective 2D and 3D images.** (A and B) Salvaged fluorescence can be collected in two configurations in a single-objective microscope by using an emission filter tilted at a small angle. (C) The top emission beam path of the 4Pi-SMS system was blocked to mimic the detection of a single-objective system. (D) Two-color 2D image of ER and microtubules in a COS-7 cell acquired using the configuration in (C). (E)

Magnified image of the boxed region in (D). (F) Two-color 3D astigmatic image of microtubules and mitochondria in a COS-7 cell acquired using the configuration in (C). The top right corner shows the data in a rainbow color table denoting z positions, and the bottom left corner shows the two-color overlay. (G) A 100-nm thick x-y slice of the boxed region in (F). (H) x-z view of the blue dashed box (200 nm wide) in (G). (J) y-z view of the orange dashed box (200 nm wide) in (G).

#### Captions for Supplementary Movies

**Movie S1. Conventional and salvaged fluorescence images of AF647 and CF660C.** The displayed raw camera frames are from two recordings of single-color microtubule samples recorded at 100 Hz shown in (fig. S4, A and D). The movie is played at 30 Hz and displayed at the same contrast as (Fig. 1B).

**Movie S2. Two-color 4Pi-SMS images of ER in a COS-7 cell.** Video rendering of the data shown in (Fig. 2, A to D; fig. S6, A to C). Two-color images of ER membrane (overexpressed GFP-Sec61β labeled with anti-GFP antibody; magenta) and ER lumen (overexpressed mCherry-KDEL labeled with anti-RFP nanobody; green).

**Movie S3. Two-color 4Pi-SMS images of mitochondria in a HeLa cell.** Video rendering of the data shown in (Fig. 2, F to I; fig. S7, A to C). Two-color images of the outer mitochondrial membrane (anti-TOM20 antibody; magenta) and mitochondrial nucleoids (anti-dsDNA antibody; green). The dataset is reconstructed from 4 optical sections with 500 nm-step sizes.

**Movie S4. Multicolor 4Pi-SMS images of the synaptonemal complex in mouse spermatocytes.** Part I: two-color images of synaptonemal complexes labeled with anti-SYCP3 antibody and anti-SYCP1 C-terminal antibody (same data as shown in fig. S8, A and B). Part II: two-color images of synaptonemal complexes labeled with anti-SYCP3 antibody and anti-SYCP1 N-terminal antibody (same data as shown in fig. S8, C and D). Part III: three-color image of synaptonemal complexes labeled with anti-SYCP3 antibody, anti-SYCP1 C-terminal antibody and anti-Lamin B antibody (same data as shown in fig. S8, E and F). Each dataset is reconstructed from 20-21 optical sections with 500-nm step sizes.

**Movie S5. Three-color 4Pi-SMS images of the Golgi apparatus in HeLa cells.** Part I: three-color imaging of *cis* Golgi (anti-GRASP65 antibody), *medial* Golgi (overexpressed ManII-GFP and labeled with anti-GFP nanobody), and *trans* Golgi (anti-p230 antibody) markers (same data as shown in Fig. 3 and fig. S9). Part II: three-color imaging of *cis* Golgi (anti-GRASP65 antibody), *cis* Golgi (anti-GM130 antibody), and *medial* Golgi (overexpressed ManII-GFP labeled with anti-GFP nanobody) (same data as shown in fig. S10). Each dataset is reconstructed from 4 optical sections with 500-nm step sizes.

**Movie S6. Two-color 4Pi-SMS images of ER-PM contact sites in COS-7 cells.** Video rendering of the data shown in (Fig. 4, A and B; fig. S11A). Two-color images of ER (overexpressed GFP-Sec61β labeled with anti-GFP antibody) and PM (labeled with WGA) in control cells (Part I), and in cells overexpressing mCherry-ORP5 (Part II) or mCherry-ESYT2 (Part III).

**Movie S7. Three-color 4Pi-SMS images of ER-PM contact sites in COS-7 cells.** Part I: three-color images of Sec61β, ORP5 and WGA (same data as shown in Fig. 4H and fig. S12G). Part II: three-color images of Sec61β, ESYT2 and WGA (same data as shown in Fig. 4J and fig. S12H).
